# Squid: Simplifying Quantitative Imaging Platform Development and Deployment

**DOI:** 10.1101/2020.12.28.424613

**Authors:** Hongquan Li, Deepak Krishnamurthy, Ethan Li, Pranav Vyas, Nibha Akireddy, Chew Chai, Manu Prakash

## Abstract

With rapid developments in microscopy methods, highly versatile, robust and affordable implementations are needed to enable rapid and wide adoption by the biological sciences community. Here we report Squid, a quantitative imaging platform with a full suite of hardware and software components and configurations for deploying facility-grade widefield microscopes with advanced features like flat field fluorescence excitation, patterned illumination and tracking microscopy, at a fraction of the cost of commercial solutions. The open and modular nature (both in hardware and in software) lowers the barrier for deployment, and importantly, simplifies development, making the system highly configurable and experiments that can run on the system easily programmable. Developed with the goal of helping translate the rapid advances in the field of microscopy and microscopy-enabled methods, including those powered by deep learning, we envision Squid will simplify roll-out of microscopy-based applications - including at point of care and in low resource settings, make adoption of new or otherwise advanced techniques easier, and significantly increase the available microscope-hours to labs.

## 1. Introduction

The past few years have witnessed tremendous developments of microscopy hardware systems, software and techniques. These developments include, to name a few, state of the art yet accessible super-resolution, smFRET, all-optical neurophysiology and light sheet systems [1–39], low-cost microscopes with a variety of applications and demonstrated value in education [40–50], tracking microscopes that open new dimensions in studying freely behaving organisms [51–54], computational microscopy that overcome conventional physical limitations by properly combining optics and computation [55–70], computational methods and software for getting the most out of acquired data using physics and/or prior information/mining underlying structure in the data through deep learning [66, 68, 71–100], as well software for visualizing the data [101–104]. On another end of the spectrum, techniques like expansion microscopy [105, 106], spatial transcriptomics [107–113], multiplex protein imaging [114–118] expand our ways in understanding biology and disease, but at the same time are calling out for significant more microscope-hours. To fully keep up with and take advantage of these developments and advances, including new applications enabled by artificial intelligence, the availability of motorized microscopes also need to be drastically expanded, with implications on cost and footprint for individual systems.

Presently most labs leveraging imaging in their research practice have access to only a handful of microscope systems at any given time due to cost, development time, and the highly-sought after time in central imaging facilities. This limits the overall microscopy hours and freedom to implement the latest advances. To best leverage the rapid pace of microscopy development while also increasing the available microscopy hours for labs, it would be desirable to have an open, modular and relatively low-cost microscopy platform that has little or no compromise in performance when compared to existing high-end solutions. Towards this, building on our previous work Octopi [44] and others, we developed Squid, a full suite of modular and open source hardware components/configurations, and software programs, to implement facility grade widefield microscopes and other application-specific imaging platforms at a fraction of the cost of commercial solutions ($500-$8k as compared to $50k-$150k). Our platform has the added benefits of compactness, advanced features such as flat field fluorescence excitation, patterned illumination, tracking microscopy etc., as well as ease of setup, maintenance and development in both hardware and software.

Up to date, more than 15 Squids have been built with a plethora of applications including live imaging inside incubators, long term tracking of motile single cells, plate readers for implementing a SARS-CoV-2 multiplex ELISA assay [119], spatial omics and bringing high-end microscopy techniques to field settings such as ocean going research cruise (R/V Kilo Moana during the HOT 317 research cruise, Dec 2019). The modularity of our approach and several other recent platforms including Planktonscope [48], UC2 [**?**], OpenFlexure Microscope [50], Flexiscope [120], smfBox [23], miCube [8], liteTIRF [2] - highlight the importance of inter-operability of components and flexibility that is provided to the research users when adopting ever expanding microscopy techniques. In the rest of this pre-print, we describe the technical features of Squid, present some characterizations as well as early results demonstrating its capabilities.

## 2. Results

### 2.1. A full suite of hardware building blocks for configuring facility grade microscopes

To simplify development and allow rapid configuration for various applications, we created a full-suite of microscope building blocks (Figure 1). We chose to use off-the-shelf components and CNC machined parts to optimize for performance, quality control and ease of assembly. By designing the blocks ground up from these components with the goal of meeting broad microscopy use needs (but not trying to meet all possible needs), we reduced the cost by a factor of 10 to 50 compared to commercial counterparts, lowering the cost and access barrier to high performance imaging systems.

**Fig. 1.**
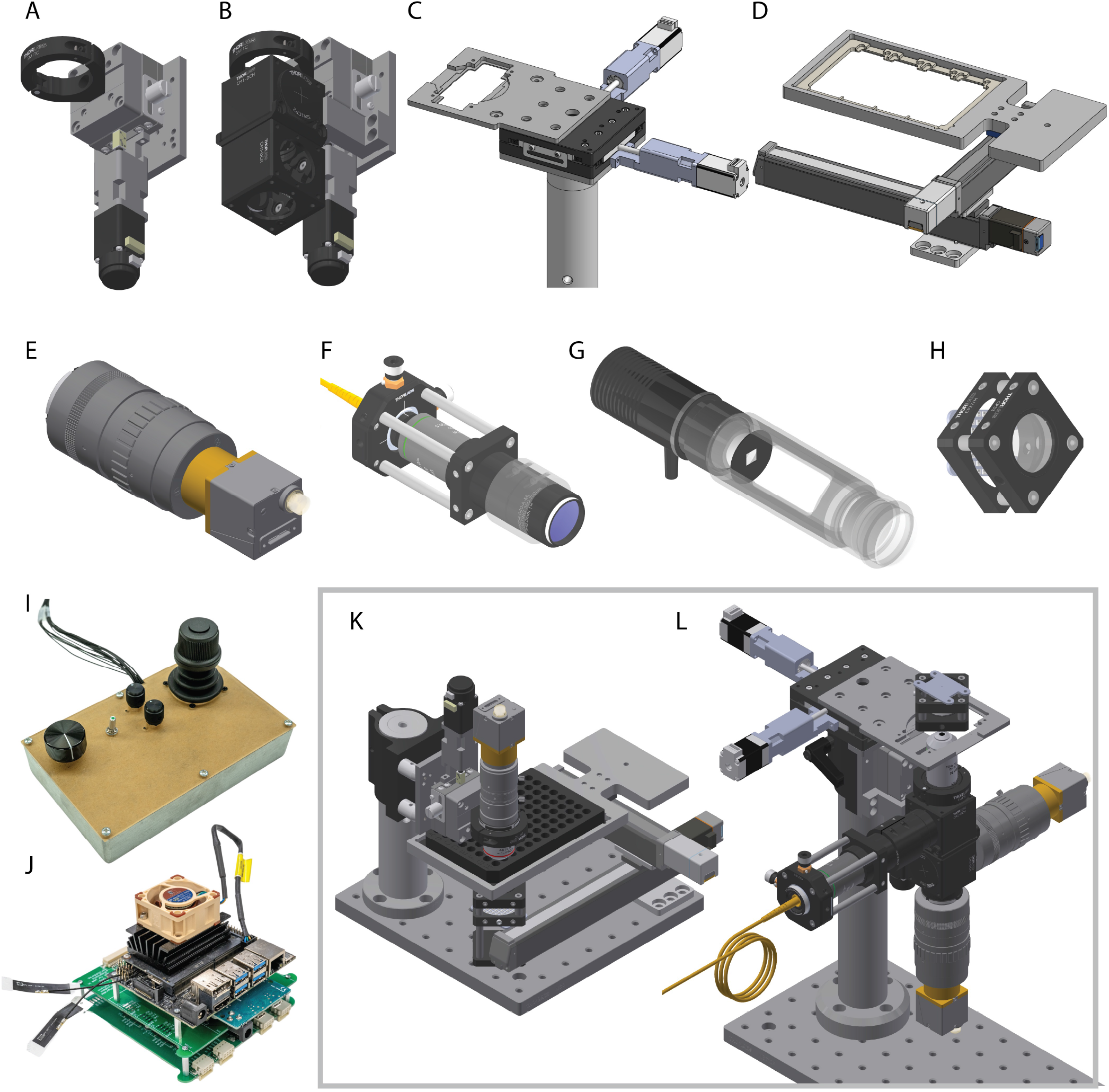
Squid hardware. (A) Motorized focus block (B) Motorized focus block with two cage cubes mounted (C) 28 mm × 28 mm motorized XY stage (D) 140 mm × 80 mm travel motorized XY stage with a well plate adapter (E) A typical image formation assembly with an industrial camera and a machine vision imaging lens (F) Flat field laser epi-illumination module (G) Flat field LED epi-illumination module (H) LED matrix trans-illuminator (I) Control panel with an analog joystick, a focusing knob, a toggle switch and two rotary potentiometers. Currently the toggle switch is used for enable tracking (when implementing tracking microscopy) and one of the potentiometers is used to adjust the XY stage max speed. (J) Driver stack (shown also a Jetson Nano for running the microscopes in place of a laptop or desktop computer) (K) One example configuration: upright microscope for reading a 96-well plate (termed Nautilus from here on-wards, see also [119]) (L) Second example configuration: multi-color flat field epifluorescence microscope with simultaneous transmitted light channel (e.g. for tracking microscopy). Various other configurations are described throughout the paper.

The microscope system can be controlled by any computer running Ubuntu (preferred) or Windows, including embedded computers like the Nvidia Jetson family. An Arduino Due is used for interfacing a control-panel, as well as for low-level control of motion, illumination, hardware camera triggers and other applications where high timing resolution and accuracy is necessary. The Arduino Due and the computer communicate through USB-based virtual COM port at 2 Mbit/s. PCB boards are implemented to allow robust integration of different electronic components. Currently a stackable, modular design is used to allow sequential development and future expansion of functionalities. Already implemented boards include a processing plane for interfacing the Arduino Due, a motion plane with 4 TMC2209 Silent Step Stick stepper motor drivers, as well as a board for the control panel and a board for interfacing the motion plane with encoders and limit switches. Notably, the driver stack can be used to support any motorized stages that are stepper motor-based. Illumination boards with 6 laser diode drivers (using TI DAC80508, a 8-channel 16 bit DAC and Wavelength Electronics laser diode drivers) and 6 LED drivers (using TI TPS92200) are currently being designed (a 8-channel DAC Arduino shield based on TI DAC8568 is also available from [121]). The board will include SMB interfaces to allow control of other commercial laser and LED engines.

While some of the modules presented in Figure 1 will be described in more detail in the following sections, Figure 2 shows some example optical configurations that can be implemented. So far we have several implementations based on Figure 2 A, B, E for a variety of applications ranging from long term imaging, plate reading, multi-color fluorescence imaging of pathology specimens to tracking microscopy, and are currently bringing online implementations based on Figure2 F and G to extend high-speed tracking to 3D.

**Fig. 2.**
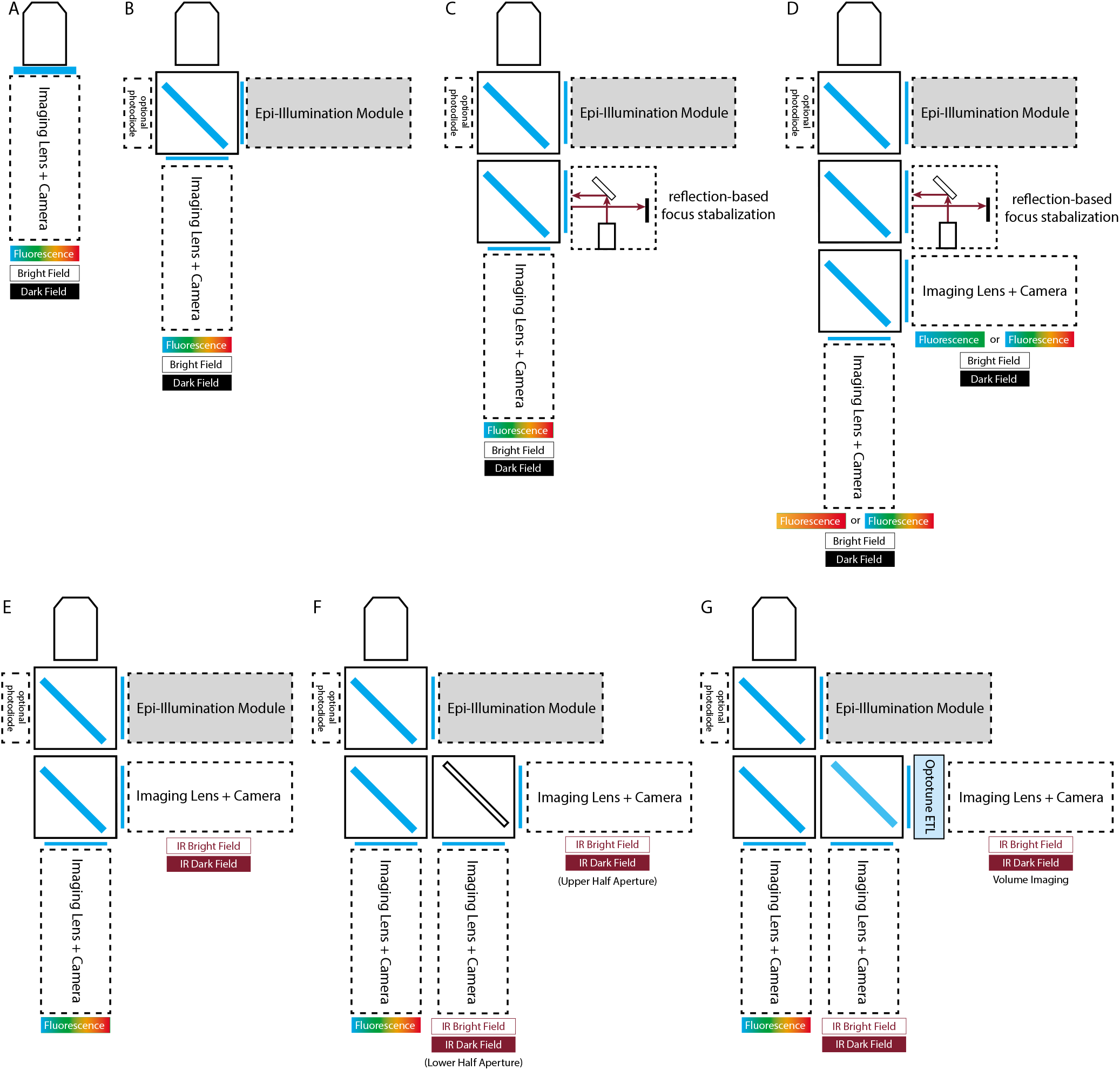
Example microscope configurations. (A) The simplest, single-camera configuration, which may be used for fluorescence with oblique angle illumination [40, 44] or waveguide illumination. (B) A single camera configuration with epifluorescence illumination. By using multiband filters and dichroics or a color camera [44], multicolor fluorescence microscopy can be achieved. The same camera is also used for non-fluorescence contrasts. (C) A single camera configuration with both epifluorescence illumination and a reflection-based focus stabilization system (e.g. [3, 7, 122, 123]). The second dichroic filter can be a 775 nm short pass dichroic. Alternatively, if bead fiducials can be introduced, the laser can be replaced with an IR LED above the sample to limit the drift to below 1 nm across all three dimensions [124–127]. (D) Another camera is added so that two or more fluorophores can be imaged at the same time [128–131]. Depending on fluorophores, filters and dichroics, either one or two lasers can be on for simultaneous multicolor fluorescence. (E) A two camera configuration with epifluorescence, where one camera is for tracking with IR illumination. (F) A three camera configuration with two cameras for both in plane and focus tracking. A mirror that divides the aperture into two cameras is used to manifest defocus as image shift. (G) A three camera configuration where one camera is used with an electrically tunable lens for volumetric imaging and focus tracking. For the arm with electrically tunable lens, a 4f system can be used to place the tunable lens at the imaged back focal plane of the objective so that there’s no magnification change when focus is swept. Alternatively, a z-splitter may be used to trade off FOV or spatial resolution with temporal resolution with one [124, 132–134] or multiple cameras [135–139]. Note all configurations can be upright, inverted, or horizontal for vertical tracking microscopy [54].

### 2.2. Python-based software

To supplement the modular hardware suite, we have developed a highly modular, object-oriented, python-based control software (with code-structuring inspiration from [3], which is based on [140]). The code is divided into drivers (for cameras, microcontroller and and other devices that may need to interface directly with the computer), core functionalities implemented as controllers (e.g. camera stream handler, threaded image saver, live controller that controls the current microscope mode and its settings, configuration manager that manages loading, updating and saving of microscope configurations; core functionalities also include application specific controllers, e.g. for tracking and for multi point acquisition), widgets that serve as interfaces between the user and the functionalities, and a top level GUI that instantiates different controllers and widgets and links the objects together. This approach to software has made it very easy to switch between microscope hardware configurations as well as further expand functionalities. Figure 3 and Supplementary Video 1 show an example of one graphical user interface.

**Fig. 3.**
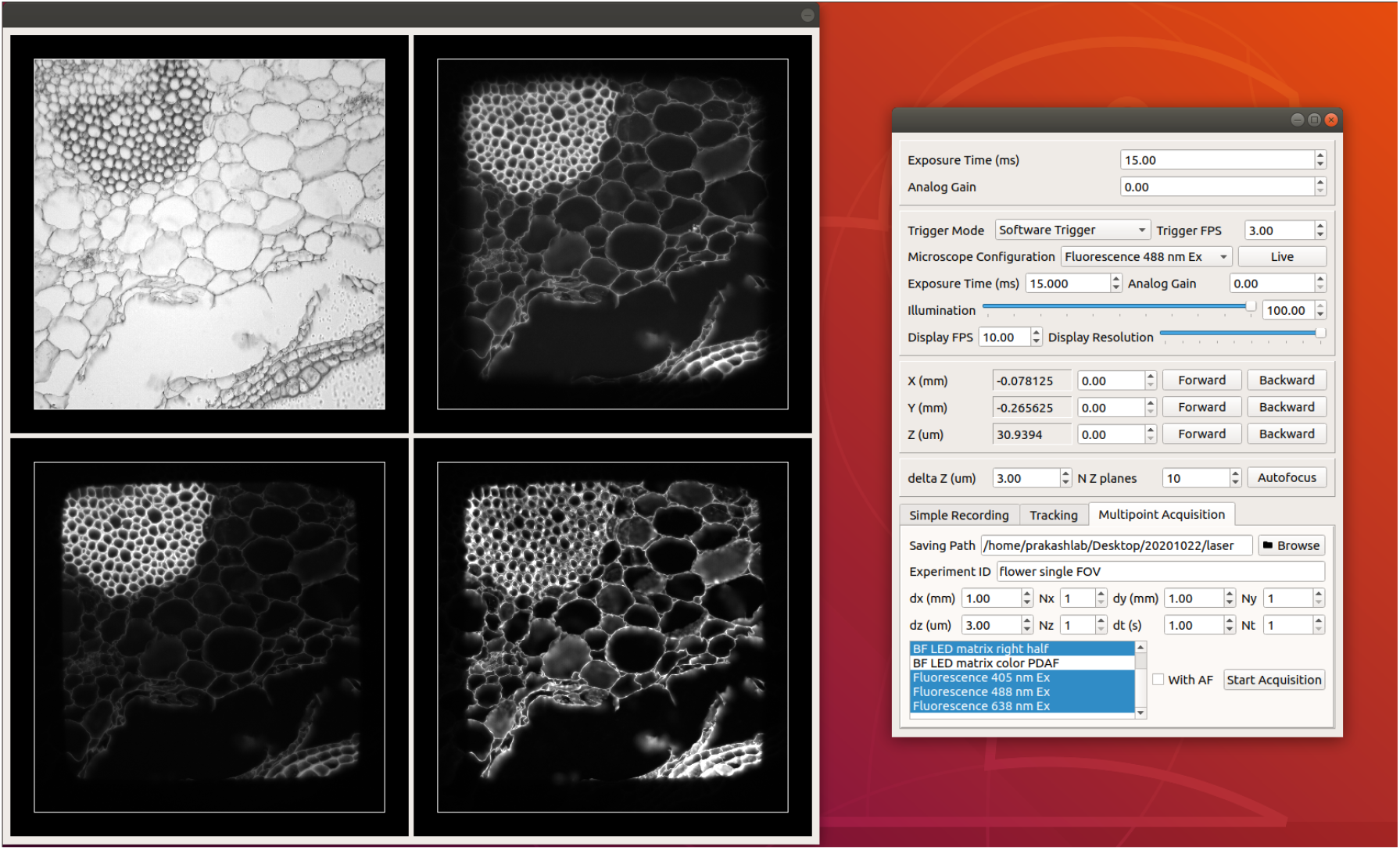
An example graphical user interface for the microscope. Using this GUI, user can set camera and illumination parameters for different configurations, move the sample, adjust focus, and control multi-point, multi-configuration acquisitions.

### 2.3. Motorized focus block and XY stage

One central piece available from the suite of building blocks is the motorized focus block (Figure 1A), which is constructed with a ball bearing linear stage with a coupled linear actuator. A piezo stack is optionally inserted between the stage and the shaft of the linear actuator to allow nm resolution focus adjustment. The use of a Thorlabs lens tube clamp for mounting an optical train (objective, imaging lens, cameras and necessary adapters for connecting them) makes it easy to quickly switch between different optical trains without taking apart the assembly, where Thorlabs SM1 lens tube and the associated adapters allow most objectives to be mounted. For more complex optical trains, the block provides direct mounting of cage cubes (Figure 1B), which decouples the objective from the rest of the optical train, allowing the objective to be moved for focus adjustment while other components remain stationary, similar to what’s in commercial microscope bodies. The focus block can be mounted either through a post mounting clamp to a 1.5” diameter post, or on a machined structure for easily and robustly integrating with longer downstream optical paths or more sophisticated optical setups.

Characterization of the motorized focus block shows sub-200 nm resolution with 1/8-microstepping and good open-loop repeatability (Supplementary Figure 1A-H). We note that when doing z-stack in open-loop, to achieve good repeatability and uniform step size allowed by the mechanics, a back-and-forth maneuver is needed to counteract mechanical backlash in the linear actuator (Figure 1I-L), which is present despite the actuator being spring-loaded by the linear stage. We also found that due to the mechanical backlash and limitation of lead screw accuracy, rotary encoders have very limited utility in this application (Figure 1A-K). To support closed-loop positioning for improved accuracy and repeatability, we’re currently integrating an optical encoder that supports 5 nm resolution and can be used for homing purpose (Supplementary Figure 2). In the characterization experiments, we also show that the focus stage can move at speed of at least 5 Hz with 180 *µ*m peak-to-peak amplitude (sinusoidal) when driven by the linear actuator and 30 Hz with 2.5 *µ*m peak-to-peak amplitude (sinusoidal) when driven by the piezo stack (limited by the travel of the piezo stack; a longer piezo stack with larger travel, e.g. Thorlabs PK2FSF1 that has 220 *µ*m free stroke, can be used if necessary). For applications where faster scanning or scanning a larger range is needed, remote focusing with a tunable lens or a higher speed stage is more suitable [141]. We also measured z-drift over more than 30 hours and found the likely temperature-correlated drift to be 1*µ*m.

A motorized XY stage with 28 mm × 28 mm travel is constructed similarly to the z-axis stage (Figure 1C). Currently we’re using a NEMA-8 linear actuator with 5 *µm* full step size, which allows max speed of about 10 mm/s. The max speed may be increased to 120 mm/s by switching to larger NEMA 11 or 14 linear actuator, reducing the spring constant of the springs in the ball bearing XY stage, increasing the drive current of the linear actuator or a combination of these. A sample holder was designed for mounting standard glass slides, cover slips and petri dishes. This module can be directly mounted on a 1.5” post, which in the inverted configuration, can be the same post used for the focus block.

For applications that requires larger travel, we have made use of linear guide rails-based, ball screw driven motorized stages (Figure 1D) and lower cost lead screw driven motorized stages [54]. We did find parallelism performance issue for the ball screw driven stage employing a NEMA 8 stepper motor and in future revisions we will use stages with cross roller bearings.

Given the unique construction of the microscope (Figure 1K-L) - including the way lenses and cameras are mounted (Figure 1E; also discussed in the next section), and that the microscope may not sit on an optical table, we set out to characterize the stability of the system and its immunity to vibrations, if any. We did so by imaging single or clusters of 83 nm diameter YG fluorescent beads (Polysciences, cat# 17150-10) with a 60x/1.49 NA objective. When the breadboard sits directly on a table, we did observe oscillation at around 60 Hz with peak to peak amplitude of around 40 nm (Figure 4 A,B). Adding 4 sorbothane feet (Thorlabs AV3) underneath the breadboard effectively suppressed the oscillation (Figure 4 B,C).

**Fig. 4.**
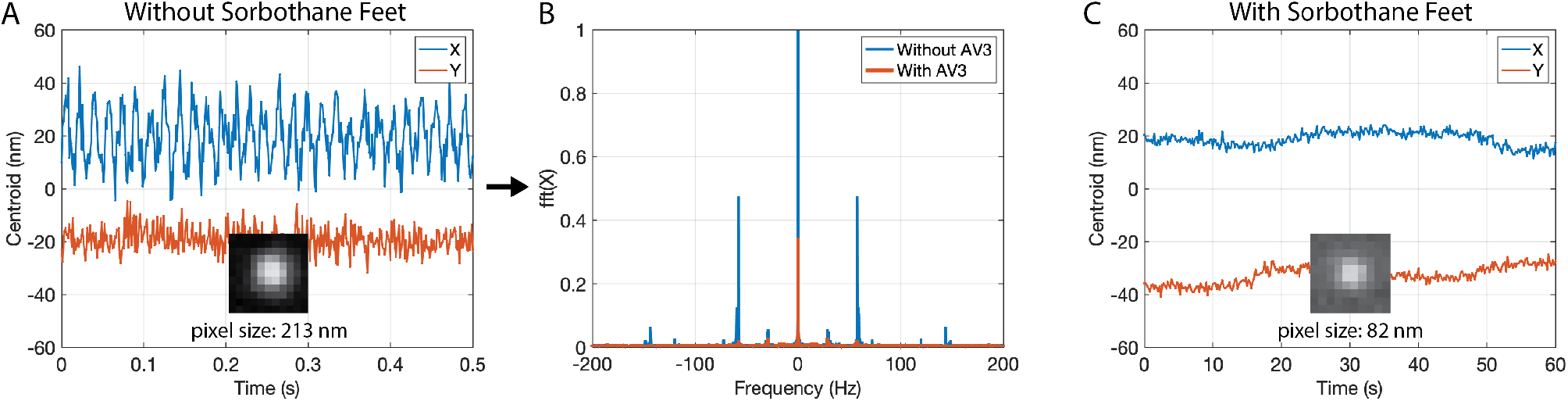
Stability of the system. (A) Time trace of the centroid of a cluster of beads (camera sensor: On Semiconductor Python 300, exposure time: 1 ms). Inset: image of the cluster of beads used for localization. (B) Fourier Transform of the time trace of the centroid (along x), showing a dominant peak at around 60 Hz when AV3 Sorbothane feet are not used. This peak and other peaks are effectively surprised after AV3 Sorbothane feet are added. (C) Time trace of the centroid of a single bead (camera sensor: Sony IMX226, exposure time: 50 ms). Inset: image of the bead used for localization). The drift over a minute in this particular measurement is within 20 nm, which is comparable to the reported drift on a Nikon Ti with perfect focus after 30 min equilibration [2].

### 2.4. Imaging Lens and Camera Assembly

Another building block extensively used in different configurations is the imaging lens + camera assembly, which consists of a machine vision lens and an industrial camera. The industrial cameras uses the latest generations of CMOS sensors with convenient USB3, GigE or MIPI interfaces. A wide range of sensors with different number of pixels, pixel size, frame rate, shutter (rolling vs global) can be selected based on applications. The smaller pixel size allows use of lens of shorter focal length (50 mm or 75 mm as compared to 165 mm - 200 mm) for image formation, making the system more compact. While commercial tube lenses at these focal length does not currently exist, high resolution (5 MP-20 MP) machine vision lenses properly corrected for different aberrations are available at relative low cost ($100-$500). To compare the performance of our imaging lens + camera assembly with its counterpart in commercial setups, we imaged the same region of cells with labeled cDNA amplicons with a f = 200 mm tube lens that’s part of a Nikon Ti2 microscope + a Prime 95B sCMOS camera (2.6 MP, 11 um pixel size) and a f = 50 mm machine vision lens (HIKROBOT MVL-HF5028M-6MP) with an industrial camera (Daheng Imaging MER-1220-32U3M) using Sony IMX226 (12 MP, 1.85 um pixel size), using the same exposure time. For the Prime 95B, an additional 1.5X magnification is used so that the objective side pixel size is similar to that of the Sony IMX226 (122 nm vs 127 nm). For the Sony IMX226, a 4f system is used to bring the back focal plane of the objective outside the microscope body so that the f = 50 mm imaging lens can be used. Figure 5 shows that an industrial CMOS camera paired with a high resolution imaging lens can deliver results as good as from the much more expensive commercial counterparts (see [19, 88, 125, 135, 142–144] for more studies comparing the use of an industrial CMOS sensor vs the use of an sCMOS or EMCCD).

**Fig. 5.**
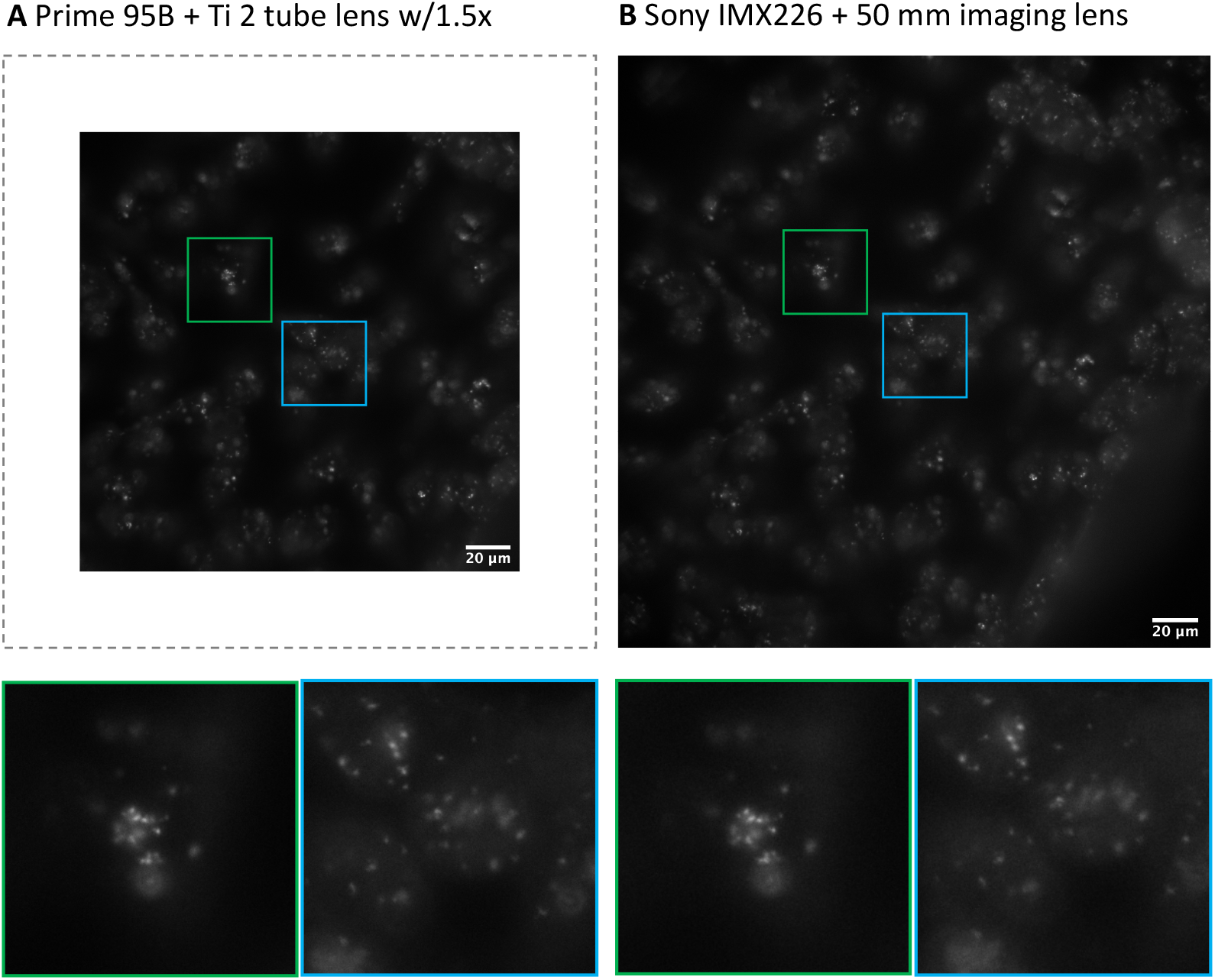
Comparing one of our image formation assembly ((HIKROBOT MVL-HF5028M-6MP + Daheng Imaging MER-1220-32U3M) with its counterpart in Nikon Ti2 with a Prime 95B camera. The contrasts are linearly adjusted for better comparison. Note that in our latest implementations we use HIKROBOT’s newer 50 mm machine vision lens (MVL-HF5024M-10MP) that has a larger aperture and better supports small pixel size.

The wide range of available industrial CMOS sensors makes it possible to choose a camera with a suitable sensor for a given imaging experiment. For example, we used a camera with an On Semiconductor Python 300 sensor supporting 860 fps with a 100x/1.25 oil objective to image three different ultrafast events in *Vorticella sp*. (Figure 6, Supplementary Video 2). The same camera and other global shutter CMOS cameras with selectable ROI (e.g. IMX250) can be used for high speed volumetric imaging or focus tracking when combined with a tunable lens. In another setup, we took advantage of the Sony IMX250MZR sensor with integrated polarizers on top of pixels to perform single shot polarization imaging (Supplementary Figure 4), which would otherwise require a more complex setup [145, 146]. This same polarization camera can be used for volumetric imaging of intrinsic density, anisotropy, and 3D orientation of cell and tissue components [66, 67].

**Fig. 6.**
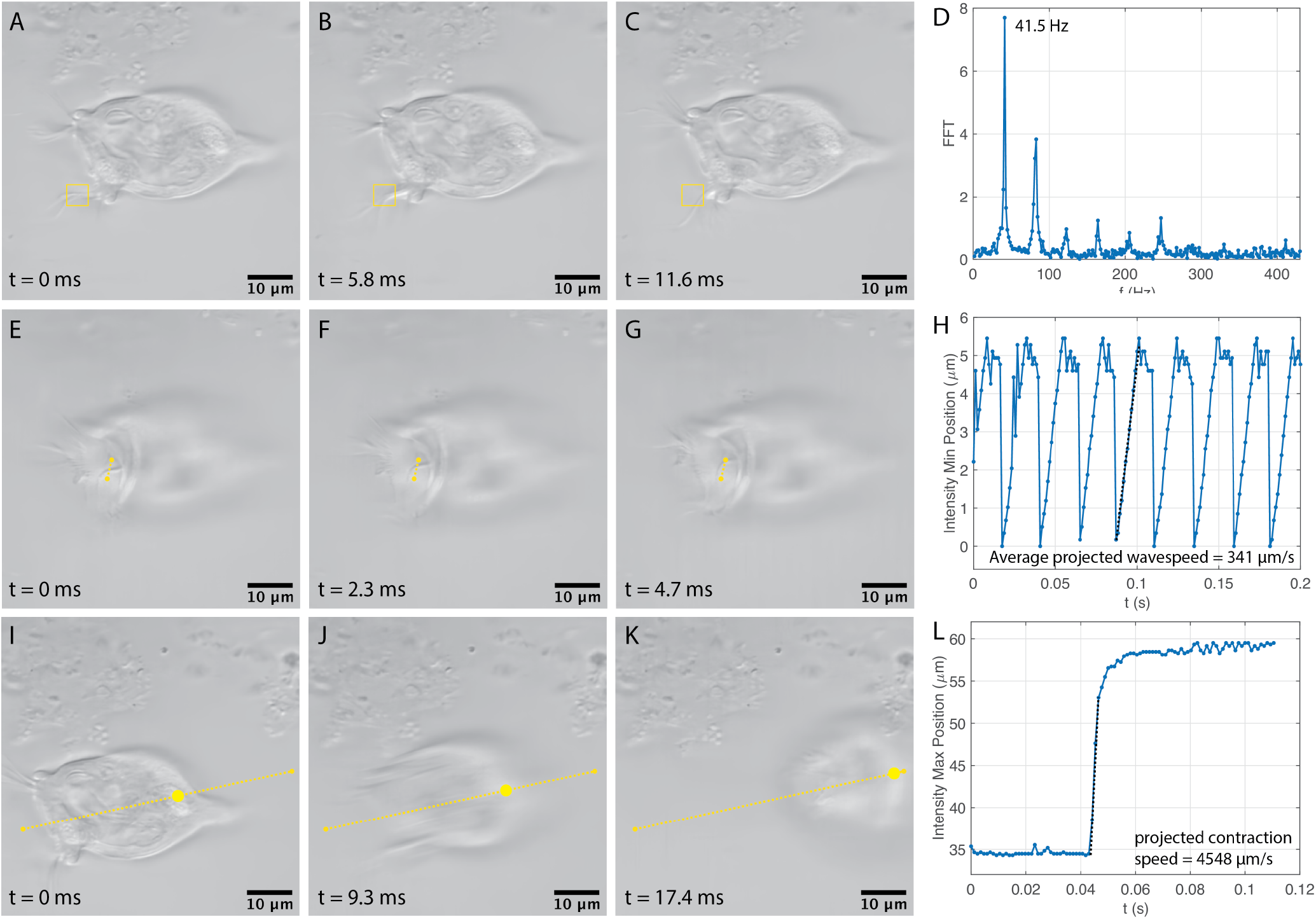
Demonstration of high speed imaging with an On Semiconductor Python 300 CMOS sensor using *Vorticella sp*. as an example. (A)-(D) Ciliary beat frequency estimation by identification of dominant frequency in the time series data (0.58s, 500 frames, 860 fps) of the minimum pixel intensity value in the selected box. The observed frequency of 41.5 Hz at 25°C is close to previously reported value of 40 Hz [147]. (E)-(H) Metachronal wave speed estimation for the collective motion of oral cilia using time series data (0.58s, 500 frames, 860 fps) from position of pixel intensity minima measured along the annotated line. The projected speed is 341 *µ*m/s *τ* = 1071 *µ*m/s, which is close to previously reported value of 1000 *µ*m/s [147, 148]. (I)-(L) Stalk myoneme contraction speed calculation from a time series (single event over 5 frames) of pixel intensity maxima position measured along the annotated line. Images shown here are denoised by FFDNet [149]. Raw images can be found in Supplementary Figure 3.

### 2.5. LED array Illuminator

To allow programmatically controlled illumination (not only intensity but also NA and angles) [57] and fast single or two-shot autofocus [150], we implemented an illuminator consisting of an 8×8 programmable LED array and a condenser with a diffuser surface (Figure 1 H, Figure 7). This illuminator and its variations (e.g. LED matrix with more LEDs, LED ring) may be used to enable techniques including quantitative phase imaging, intensity diffraction tomography and Fourier ptychographic microscopy. Other illumination modules that have been used in these applications can also be adopted [57, 61, 69, 151, 152]. We are further incorporating an LCD-TFT display for finer control of the illumination, which can be used in recently developed techniques in quantitative phase imaging [59, 67].

**Fig. 7.**
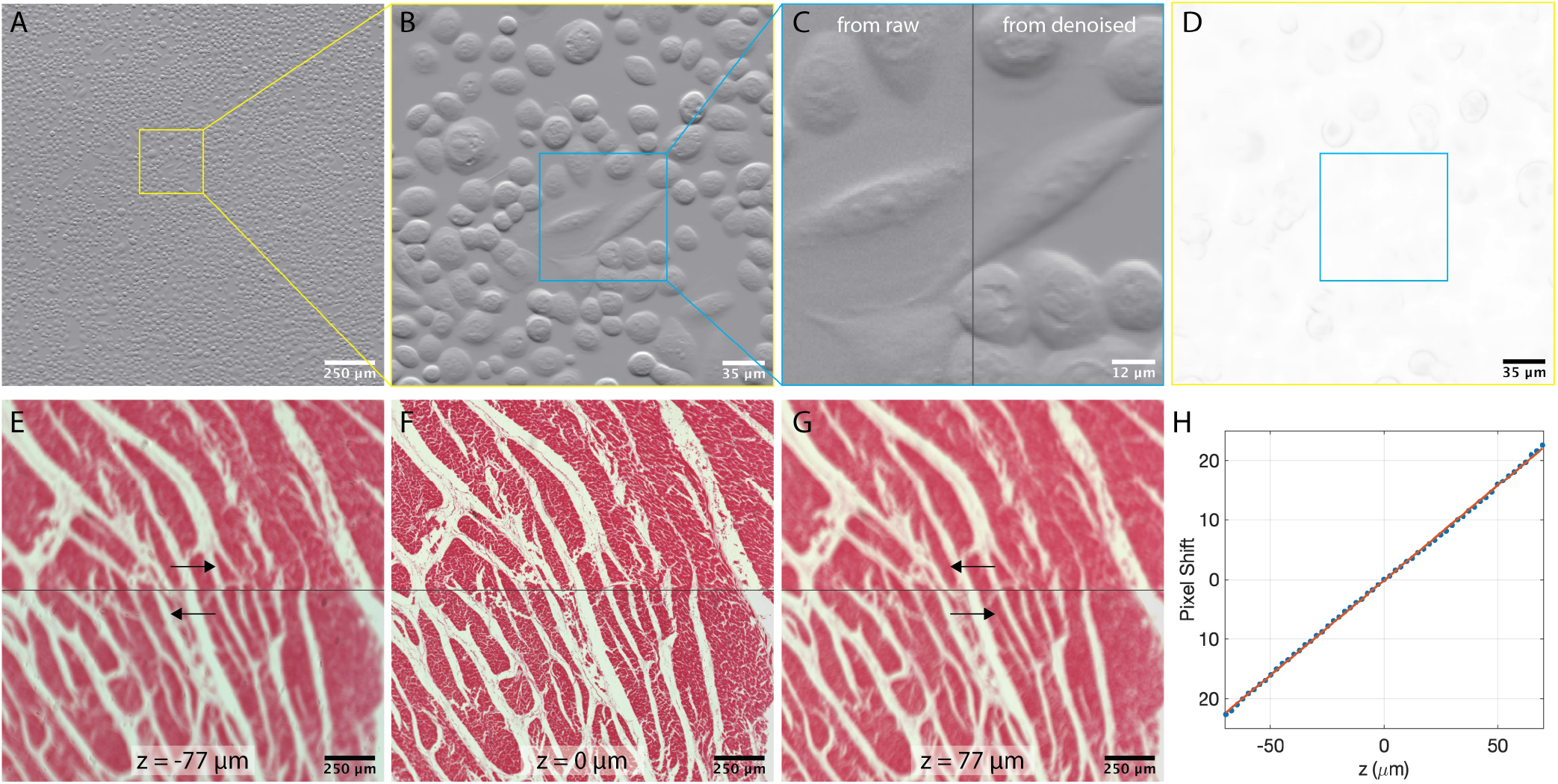
Imaging with a LED matrix illuminator. (A-C) Differential Phase Contrast of culture of human conjunctival epithelial cells. Compared to bright field (D), the contrast is significantly improved. (C) also compares the DPC result from raw images and images denoised with FFDNet. (E-H) Fast two-frame autofocus. The amount of defocus (including the sign) can be calculated from the pixel shift of a pair of images with opposite halves of the LED turned on [150].

### 2.6. Multicolor Epifluorescence Illumination and Patterned Illumination

Multicolor flat-field epifluorescence illumination is desirable in many applications. To make this feature more readily accessible, building on top of others’ work [18, 153, 154], we have developed an easy-to-integrate flat field launch using square core multimode fiber (Figure 1 F and Figure 8 A inset) and a low-cost (about $1000), compact and alignment-free multi-color laser engine (Figure 8 A). Because the magnified image of square fiber core (formed by the 80x objective) is imaged into the infinity space of the imaging objective, translation of the objective or switching objectives would not affect the image of the fiber (the illumination profile) in the object plane. By adding a translation module, TIRF and HILO (highly inclined and laminated optical sheet) [155] illumination can be achieved.

**Fig. 8.**
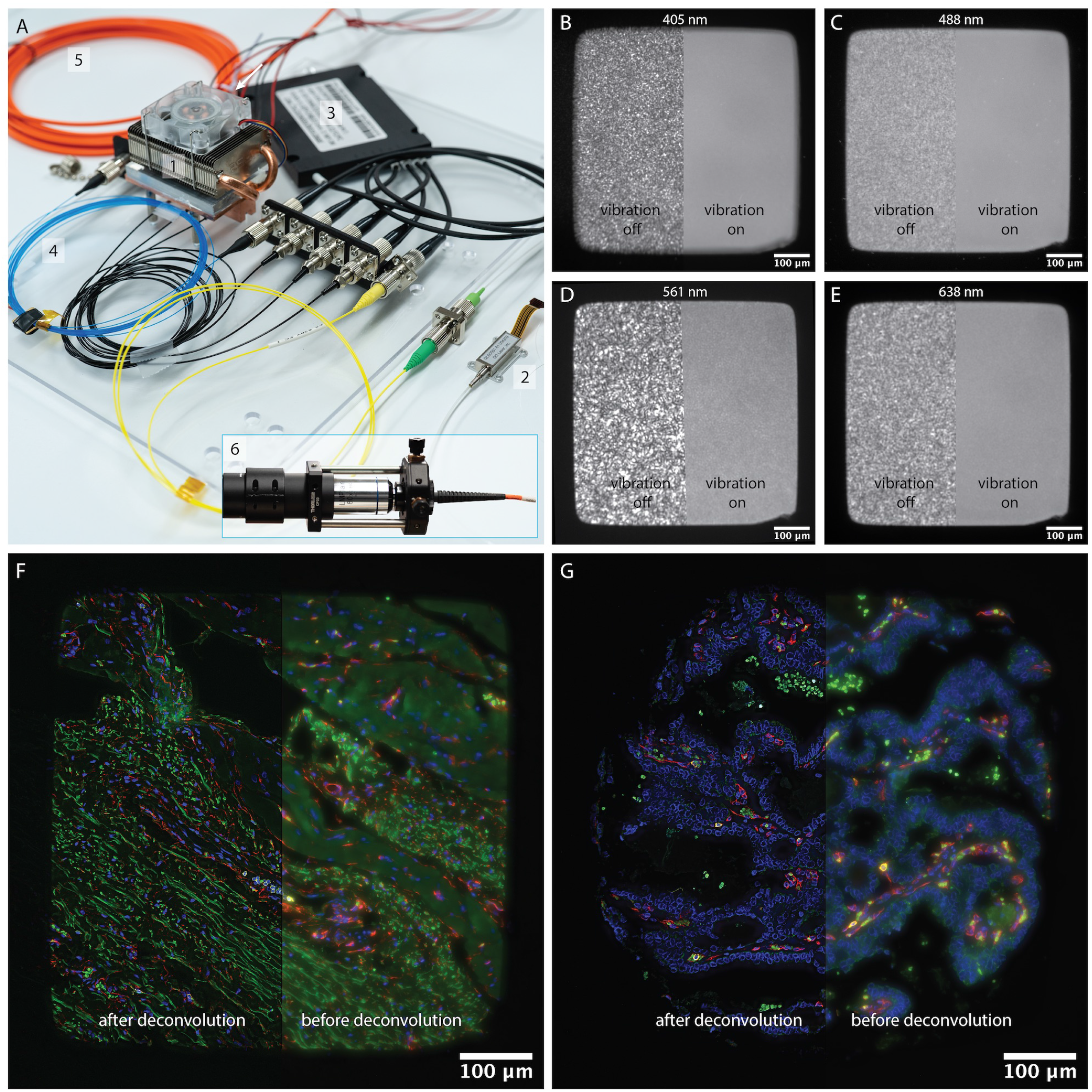
Flat field multicolor laser epifluorescence illumination. (A) Photo of the laser engine and flat field launch. 1: three multimode fiber pigtailed laser diodes (405 nm, 488 nm and 638 nm) mounted in a custom machined copper block attached to a Jetson Nano heatsink, 2: 561 nm fiber pigtailed laser module from QDLaser, 3: fiber combiner, 4: fiber-shaking speckle reducer, 5: 150 *µ*m square core fiber, 6: flat field launch consisting of an 80x metallurgical objective and a 40 mm Hastings triplet. If square core fibers with larger core sizes are used, an objective with lower magnification or an achromatic lens may be used. (B-E) Illumination profile measured with highly concentrated thin fluorescent film (Materials and Methods) (F-G) Max intensity projection of z-stacks (17 planes, 1.5 *µ*m spacing) of a CODEX-processed tonsil sample and tissue core (blue channel: nucleus stained by Hoechst, green channel: autofluorescence, yellow channel: CD45 stained by Cy3, red channel: CD34 stained by Cy5). Deconvolution is done with an unmatched back projector [77]

Using 105 *µ*m multimode fiber pigtailed laser diodes in our laser engine and a quad band filter set, we are able to get 43.3 mW of 638 nm, 12.4 mW of 488 nm and 10.3 mW of 405 nm excitation at the object plane of a Nikon 20x/0.75 apochromatic objective (for the second laser engine we constructed, we’re able to get 60.6 mW of 638 nm, 23.6 mW of 488 nm and 20.3 mW of 405 nm after a 4x achromatic objective, and 58.9 mW of 638 nm, 19.5 mW of 488 nm and 12.8 mW of 405 nm after a 10x achromatic objective). To increase the laser power, besides choosing laser diodes with higher output power (e.g. a 700 mW laser diode at 638 nm is readily available from the same supplier), additional laser diodes can also be added. As for 561 nm, we have evaluated a compact, frequency doubled quantum dot laser from QDLaser with up to 15 mW output in single mode fiber (versions with 25 mW output using a multimode fiber and 50 mW free space output also available), which can be directly modulated up to 100 MHz - an advantage over diode pump solid state (DPSS) lasers frequently used at this wavelength. We have also recently identified a lower cost ($ 1200), 150 mW fiber-coupled Nd:YVO4 DPSS laser with PPLN intracavity frequency doubling and will be evaluating it soon. Currently, speckle reduction is implemented with shaking an inserted section of multimode fiber with a coin vibrator (15000 rpm), which performs well (Figure 8 B-E) but can also be further improved, especially for the more coherent 561 nm line. As an example application, we imaged two CODEX-processed samples (Figure 8 F-G, Supplementary Video 3), where z-stacks (17 planes, 1.5 *µ*m spacing) were taken and the results are deconvolved with an unmatched back projector [77].

We note that using the laser engine and flat field launch, HiLo microscopy [156, 157] can be readily applied, although the requirement of speckle being fully developed for the speckle image will limit the imaging speed for multicolor acquisition as the laser diodes are currently switched on/off directly. To break this limit without requiring expensive electrical or acoustic modulators (or requiring the lasers to be free-space coupled so that mechanical shutters can be used), additional dichroic beam splitter and detection paths can be added so that multiple lasers can remain on at the same time. For applications requiring single spatial mode (e.g. light sheet generation), single mode pigtailed laser diodes or modules and single mode fiber combiners are also readily available.

As an alternative to laser-based illumination, we have also implemented LED-based flat field illumination (Figure 9). Because of the large étendue of LED source, only a small fraction of optical power will be delivered to the object plane. Using a Thorlabs M405D2 at 1.2A driving current, which results in > 1.5 W emitted optical power, we’re able to get 64 mW at the object plane after a 10x/0.25 NA objective, for illuminated area of 1.65*mm* × 1.65*mm*. Multiple LEDs can be combined with light delivered to the microscope in free space or through a liquid light guide (many customizable off-the-shelf solutions are already available, e.g. from Prizmatix and bluebox optics). Higher power LEDs (>10W optical power at the LED) widely used in 3D printing and projectors are available at low cost, and can be used if appropriate heat sink is implemented.

**Fig. 9.**
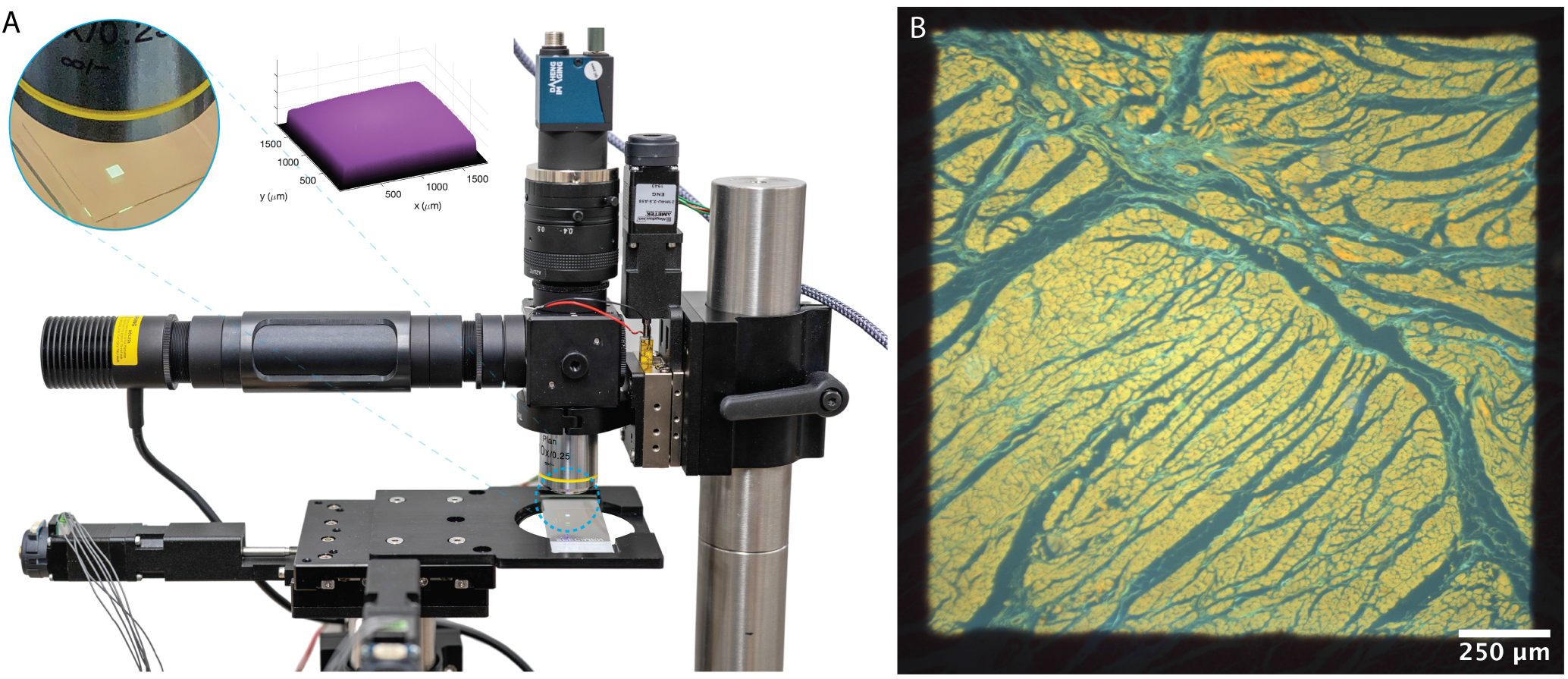
Flat field LED epifluorescence illumination. (A) Photo of the setup. The LED is collimated with an aspheric condenser (Thorlabs ACL25416U-A). A 75 mm achromatic lens (Thorlabs AC254-075-AB) is inserted in the optical train in the 4f configuration with the objective lens. A square aperture is placed at the plane conjugate to the object plane as a field stop. Insets show photo of the square illuminated spot and the measured profile. (B) Fluorescence from a prepared tissue section with 405 nm excitation imaged with a 10x/0.25 objective and a color camera.

Further more, using the configuration in Figure 2B, we demonstrated patterned illumination with a DLP projector (Figure 10), which, along with its LCoS alternatives, may be used to perform targeted photo-stimulation [24–27, 158] and structured illumination for optical sectioning and/or super resolution [19, 21, 25, 52, 137, 159, 160]. Alternatively, lasers may be used in place of the LED for higher efficiency or for coherent SIM [20, 161–164]. In applications where laser lines at 450 nm, 520 nm and 640 nm suffice, RGB laser projectors may be used for compact integration.

**Fig. 10.**
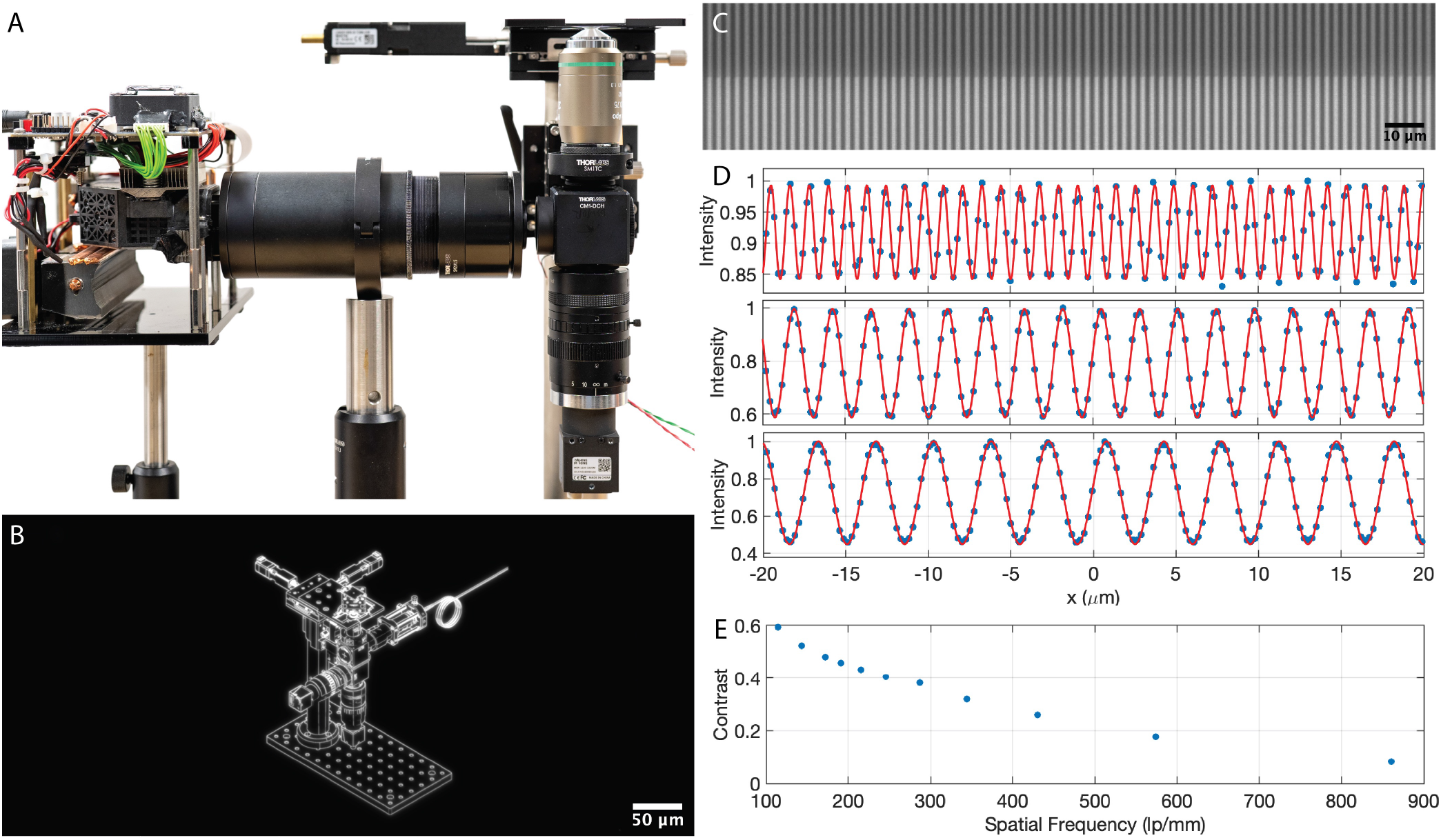
Patterned illumination with a DLP projector. (A) Photo of the setup. The projection lens is replaced with a f = 180 mm tube lens (Olympus SWTLU-C). A 20x/0.75NA objective is used. A mirror at the object plane is used for the pattern measurement (B) An example of projected pattern. (C) Projected red and green stripes imaged with a monochrome sensor. (D) Measured (blue dots) and fitted (red lines) illumination profile with different stripe spatial frequencies). (E) Achieved contrast as a function of spatial frequency in the object plane.

### 2.7. Tracking Microscopy

Conventional, fixed field-of-view (FOV) microscopy techniques are not suitable for studying live, highly motile, suspended organisms capable of complex and fast movements. To image these cells during unrestrained behavior, one is forced to sacrifice optical resolution to gain a larger FOV. Tracking microscopy is a valuable method that solves this challenge by continuously re-positioning the sample stage or imaging system to keep the tracked organism within the microscope FOV. This allows one to obtain sub-micron resolution images of an organism while allowing unrestrained behavior over scales much larger than the organism’s size [51–54, 165]. By incorporating software elements we had previously developed [54], we imparted our Squid’s with tracking capabilities. Since the software was written in python, it is straightforward for us to integrate state-of-the-art deep learning-based computer vision trackers [166] to allow fast, robust tracking even in complex and crowded environments.

We demonstrate tracking microscopy with Squid by measuring tracks of organisms over a wide range of sizes (10 *µm* to 1000*µm*) and diverse behaviors. In particular, we tracked small flagellates such as *Chlamydomonas*, fast-swimming ciliates including *T. pyriformis, Didinium sp*. and the giant ciliate *S. ambiguum* (Figure 11, Supplementary Video 4). In all cases across diverse sizes, shapes and behaviors of the organisms, we demonstrate successful tracking and obtained trajectories at high spatio-temporal resolution while concurrently obtaining high-resolution dark-field images. While dark-field is the imaging mode shown here, it is possible to combine tracking with any of the microscope configurations described in previous sections.

**Fig. 11.**
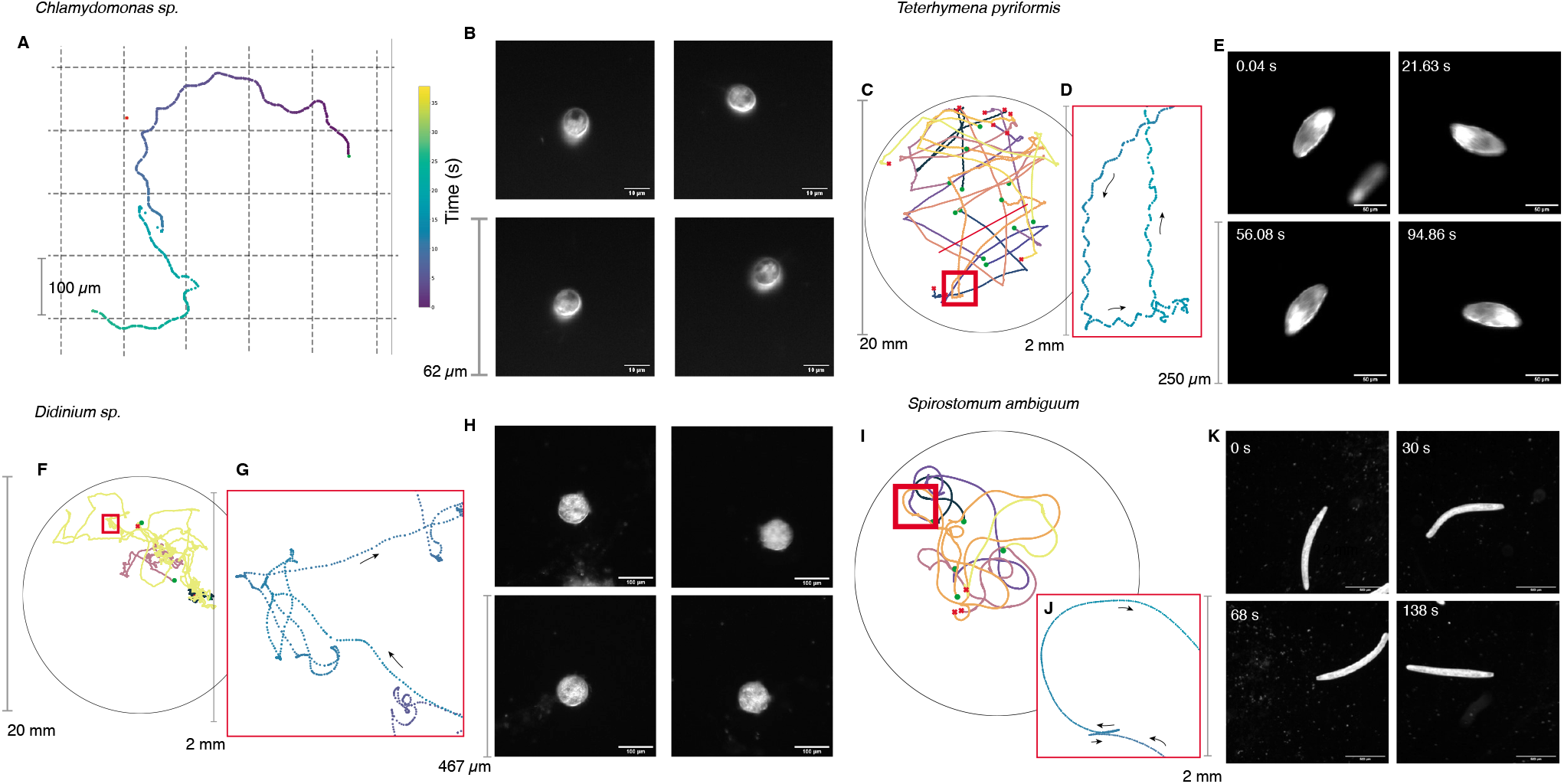
Demonstration of tracking microscopy using Squid for a wide range of organisms spanning *Chlamydomonas sp*. (size 10 *µm*), *T. pyriformis* (100 *µm*), *Didinium sp*. (300 *µm*) and *S. ambiguum* (1000 *µm*). (A) Measured track of *Chlamydomonas sp*. and (C), (F) and (I) show the measured tracks for single organisms over the scale of the chamber (diameter 20 *mm*) for *T. pyriformis* (organisms, tracks *n* = 24), *Didinium sp*. (organisms, tracks *n* = 6) and *S. ambiguum* (organisms, tracks *n* = 10), respectively. (D), (G) and (J) show zoomed-in track segments over a 2 *mm* scale, demonstrating the fine-resolution made possible by the combination of software+hardware tracking. (B), (E), (H) and (K) Snapshots of Darkfield images that show the organism being tracked with micron-scale resolution.

We further characterized the tracking performance in the experiments by calculating the distribution of tracking errors over thousands of images. The distributions (Figure 12 B-E) show that even during complex, fast swimming behaviors the tracking error remains well within the microscope FOV indicating successful tracking. Further, we demonstrate the robustness of the tracking system in complex, crowded environments by tracking a single *T. pyriformis* cell in a dense culture. Figure 12 F, G and H show the remarkable performance of the combination of both hardware and software tracking systems that are able to follow the given cell of interest in a highly-crowded environment despite multiple crossing and occlusions by cells of similar size, shape and swimming behaviors.

**Fig. 12.**
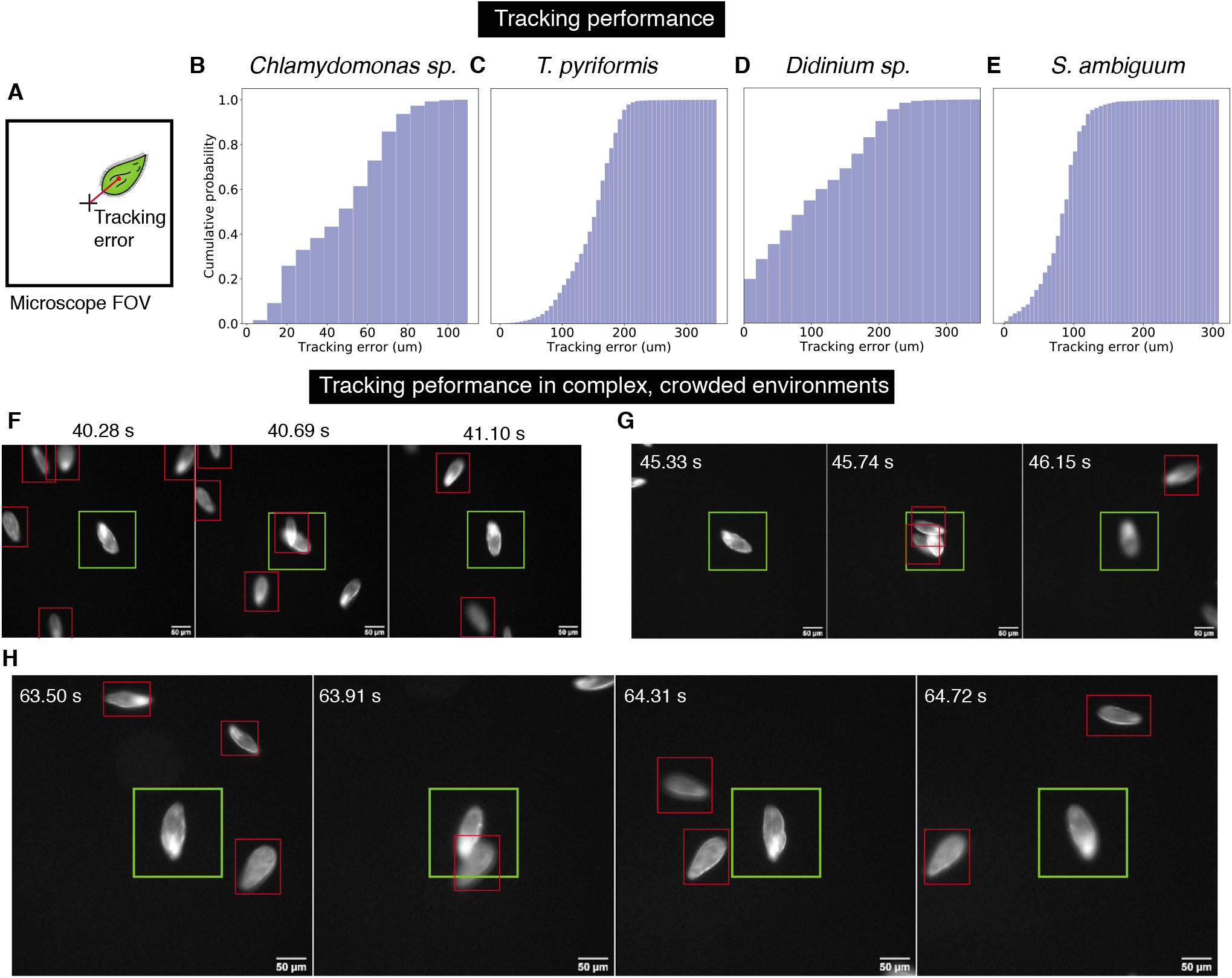
Preliminary tracking microscopy performance characterization. (A) The tracking error is used as a metric for quantifying tracking performance. (B)-(E) Cumulative probabilities of tracking error for *Chlamydomonas sp*. (calculated over 1893 images), *T. pyriformis* (calculated over 13331 images), *Didinium sp*. (calculated over images 18427 images) and *S. ambiguum* (calculated over 10730 images), respectively. (F)-(H). We used a dense culture of *T. pyriformis* to demonstrate robust software+hardware tracking in complex, crowded environments with many similar objects. The cell being tracked is shown within the green box while other cells are shown within red boxes. The snapshots in (F), (G), (H) demonstrate that the software+hardware tracking is able to keep tracking the cell of interest even during multiple cell-cell crossings as well as rapid changes in cell behaviors (not shown).

## 3. Discussion

In this preprint we described Squid - a modular platform for widefield research grade microscopes that are affordable - and present some of its technical capabilities and flexibility by demonstrating a wide variety of configurations. We also document the construction on our website at https://www.squid-imaging.org. With its high modularity, components of its hardware and software may be either built/used as is or integrated into existing solutions. While we provide a full suite of modules for a wide range of applications, users can also take advantage of other open source modules or commercially available products (motorized stages, lasers, LED engines etc.). The presented mechanical construction of Squid assumes use of compact and light weight industrial CMOS cameras, which have satisfactory performances in many applications [19, 125, 135, 137, 142–144] and can be further enhanced by generic denoising [149] or automatic correction of CMOS-related noises [88]. In applications where bulkier EMCCD or sCMOS camera are desirable or must be used (for example when the speed of the CMOS rolling shutter needs to be controlled to synchronize camera exposure with light sheet movement [30, 31, 167, 168]), additional CNC machined parts can be designed to provide needed alignment aid and mechanical support.

Squid provides a framework and the set of components for users to implement microscopes that can be tailored to specific applications with reduced budget, effort and turnaround time. With starting cost of less than $2000 (Supplementary Table 1), which includes full motorization and may be further reduced through optimization, the microscope can deliver functionalities and performances of commercial options that are 10 to 50 times more expensive. The developed motorized XY ($500) and Z stages ($300-$700 depending on encoder option), fiber coupled laser engine ($1000 for 405 nm + 488 nm + 638 nm, starting at $2000 for 405 nm + 488 nm + 561 nm + 638 nm) and tested lens and camera options (starting at $420 for a 12 MP CMOS camera and a 10 MP rated f=50 mm lens) also provide considerable cost reductions for home-builders. Beyond cost saving, high performance and versatility that have been described, Squid is designed to be easy to assemble - even the most sophisticated implementation reported in this preprint can be put together from individual parts and brought online within a single day. The constructed microscope has a small footprint, does not require an optical table to operate, and can be easily transported not only within labs but also to field sites. The open nature and standard interfaces also simplifies future expansion. Because the user has full control of the Arduino-based firmware and Python-based software, it’s straightforward to further customize the system and include additional functionalities. These attributes of Squid make it an ideal candidate for greatly increasing the accessibility of high performance research microscope as well as for rapid dissemination and adoption of wide ranging new developments in microscopy and microscopy-based techniques and otherwise advanced features that would require additional premiums in commercial solutions.

## Materials and Methods

### Microscope Hardware Components

While 3D printing has been widely taken advantage of in democratizing open source tools [47, 49, 50], for custom parts we deliberately choose to use CNC machining, which often requires parts to be ordered from vendors but offers exceptional mechanical properties, tight tolerances at price point not much more expensive than 3D printing, esp at qty greater than 5. We use Thorlabs cage and lens tube systems so that users can take advantage of their wide selection of off-the-shelf components and build the microscope with as little need in alignment as possible.

Bill of Materials (Supplementary Table 1-3) and 3D CAD files can be accessed on our website (https://www.squid-imaging.org).

### Software

The firmware that currently runs on an Arduino Due is developed with the Arduino IDE, using libraries including TMCStepper, AccelStepper, DueTimer and Adafruit_DotStar. In a future version, stepper control will be moved to hardware using stepper controller ICs (Trinamic TMC4361A). The software that runs on the host computer is written in Python and uses libraries including pyqt5/pyside2, pyqtgraph and opencv. A microcontroller class is implemented in microcontroller.py to handle communications with the Arduino Due. Controllers are implemented in core.py. Widgets that wrap around the controllers for the graphical user interface are implemented in widgets.py. Different objects that together constitute the control program are instantiated and connected in gui.py. Software (firmware included) can be found at https://github.com/hongquanli/octopi-research.

### Flat field laser illumination profile measurements

The illumination profile at the sample is measured using highly concentrated fluorescent dye samples [169], as previously described [170]. For 488 nm, 561 nm and 638 nm, we used Fluorescein Sodium Salt (FSS) (Sigma Aldrich 46970, for 488 nm excitation) and Acid Blue 9 (AB) (Tokyo Chemical Industry America, 3844-45-9, for 561 nm and 638 nm excitation). 0.9g of FSS and 0.7g of AB were dissolved in 1 ml of deionised water each to prepare separate stocks in eppendorf tubes. The eppendorfs were first vortexed and then left in an ultrasonic water bath overnight to let the smaller particles of the dyes dissolve completely. The next day, a 40 *µ*L droplet was placed on a 25 mm by 40 mm coverslip and then gently covered with a 25 mm by 25 mm coverslip. The edges of the coverslip were sealed with nailpolish resin to prevent any evaporation of the solvent. For 405 nm, we used undiluted 10 mg/ml Hochest 33342 solution in water (Invitrogen H3570).

### Patterned illumination with a DLP projector

For pattern projection we used an EKB DLP E4750RGBLC Light Control evaluation module (DPM-E4750RGBHBLCOX), with an ND = 6 absorptive filter (Thorlabs NE60A-A), and a 50:50 beam splitter (Chroma 21014-UF2). For projection with certain wavelengths, we will be using a variant from EKB with fiberlight guide coupling (EKB DPM-E4750LCFCSOXLLG).

### CODEX samples

The tonsil samples were obtained from Stanford Pathology and were fully de-identified, thus the study was exempt from ethics approval (no human subjects research). The TMA was obtained from another study [171].

### 3D Deconvolution for CODEX samples

Deconvolution was done using MATLAB programs provided with [77]. Point spread functions were generated using FIJI plugin PSF Generator using the Richard & Wolf 3D Optical Model [172].

### High speed imaging experiments

*Vorticella sp*. samples were collected from debris clusters growing along with the *Didinium sp*. cultures obtained from Carolina biological supply for the tracking experiments. The clusters were isolated from the debris using pipetting and then enclosed into a coverslip chamber (20 mm diameter, 0.12 mm depth) constructed using a coverslip spacer (Grace Biolabs).

### Tracking experiments

All tracking experiments were done in chambers 20 mm in diameter and 0.12 mm deep constructed using cover slip spacers (Grace Biolabs). Organisms were obtained from Carolina biological supply and were stored at room temperature (22 C) under white light for 3 days after shipping. Cultures were diluted in spring water 30 mins prior to tracking experiments. Tracking experiments were performed using a gaming laptop (MSI GE65 9SF Raider-051 with Intel i7-9750H CPU, Nvidia RTX 2070 GPU, 2 8 GB 2666 MHz DDR4 RAM and a Samsung 1TB 970 EVO Plus NVMe M.2 SSD). The Graphics card allow use of fast neural-network based trackers such as DaSiamRPN [173], which allowed tracking at 200 Hz.

## Acknowledgements

We thank all members of Prakash Lab for feedback throughout this work. We thank John Hickey, David McIlwain and Christian Schuerch for providing CODEX samples and helpful discussions. We thank Xiao Wang for providing the STARMap sample. We thank Chunzi Liu for providing the human conjunctival epithelial cell culture. We thank Windell H. Oskay (Evil Mad Scientist Lab) for providing CNC machining suggestions. We thank QDLaser for letting us evaluate their frequency doubled quantum dot laser module. We thank early Squid platform adopters, including Mireille Kamariza, Christina Hueschen, Eva de la Serna, Kiara Cui, Leeya Engel, Chunzi Liu, Angela Zhang, Feriel Melaine, Rishabh Shetty, Tal Sneh, Scott Coyle, Eliott Flaum, Adam Larson, Shalin Mehta’s group at CZ Biohub, Grace Zhong, Samhita Banvar, Anton Molina and Guillermina Ramirez-San Juan. The project was supported in part by financial support to from Tool Foundry through its accelerator program, Moore Foundation, Stanford Institute for Immunity, Transplantation and Infection and Bill & Melinda Gates Foundation. H.L. was supported by a Bio-X SIGF Fellowship. D.K. was supported by a Bio-X Bowes Fellowship. E.L. was supported by a NDSEG Fellowship. P.V. was supported by a Bio-X SIGF Fellowship. C.C. was supported by a NSF GRFP and Stanford Graduate Fellowship. M.P. acknowledges support from NSF Career Award, Moore Foundation, HHMI Faculty Fellows program, NSF CCC (DBI-1548297) program and CZ BioHub Investigators program.

## Author Contributions

H.L. and M.P. designed the research. H.L. designed and implemented the microscope hardware and main software. D.K. and H.L. developed the tracking software. E.L., H.L. and N.A. designed and implemented the electronics. H.L., D.K., P.V., C.C. and M.P. performed experiments. H.L. wrote the manuscript with contributions from all authors.

---

**Supplementary Figure 1.**
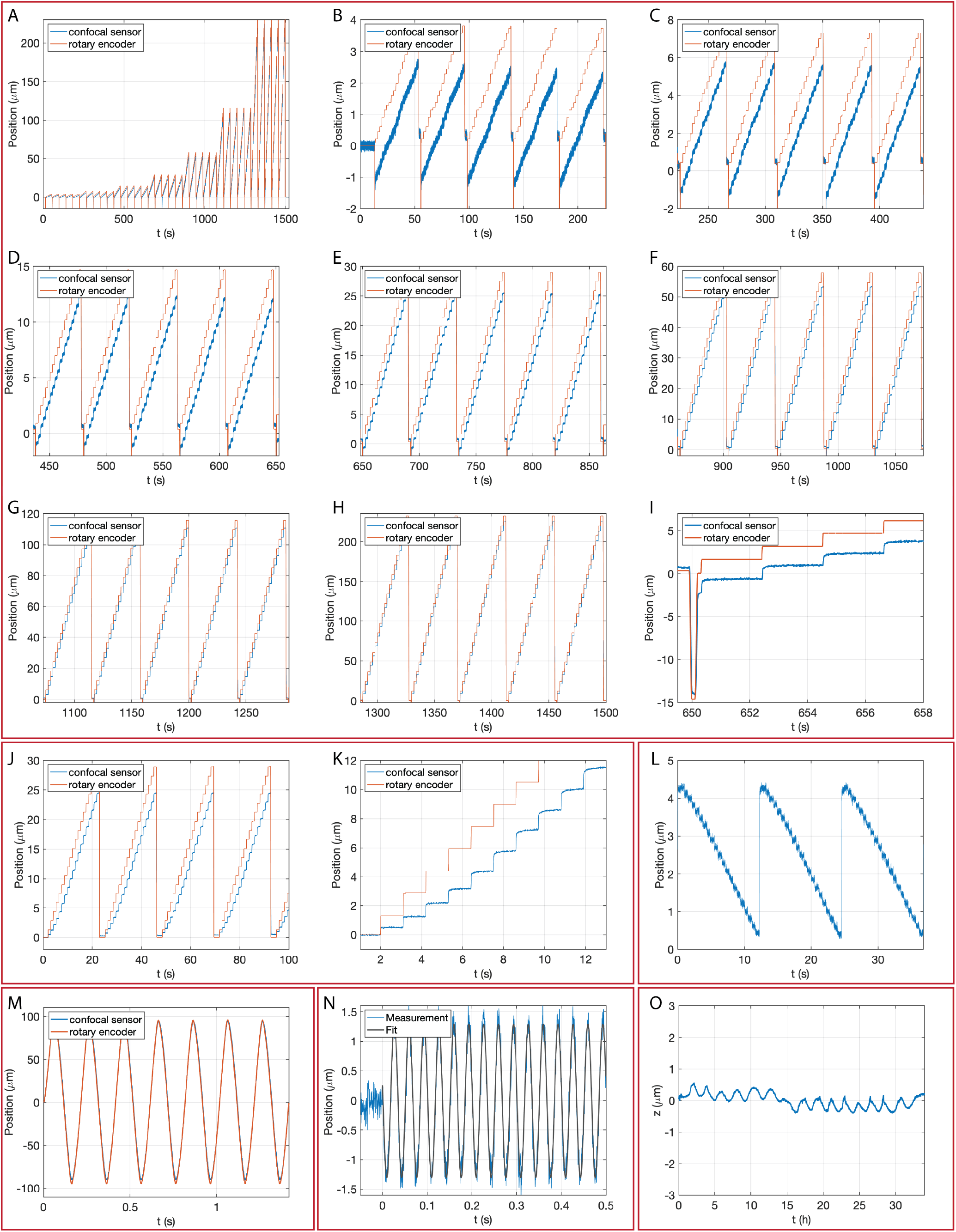
Focus stage characterization with a chromatic confocal sensor. (A) 20-step z-scan with step size of 1, 2, 4, 8, 16, 32, 64 microsteps (with 8-microstepping), corresponding to nominal step size of 188 nm, 376 nm, 752 nm, 1.5 *µ*m, 3 *µ*m, 6 *µ*m, 12 *µ*m. (B-H) Zoomed-in view of (A). (I) Zoomed-in view of the “back-and-forth” maneuver (80 microsteps) to reduce the effect of mechanical backlash. (J-K) When the “back-and-forth” maneuver is not performed, unidirectional repeatability is worse and the first few steps are smaller. (L) The step size is relatively uniform when the stage is actuated by the piezo stack, suggesting that the mechanical backlash is in the linear actuator rather than the ball bearing stage. (M) Scanning with the linear actuator at 5 Hz with peak-to-peak amplitude of 180*µ*m (N) Scanning with the piezo stack at 30 Hz with peak-to-peak amplitude of 2.5*µ*m (O) Z-drift measurement over 30 hours.

**Supplementary Figure 2.**
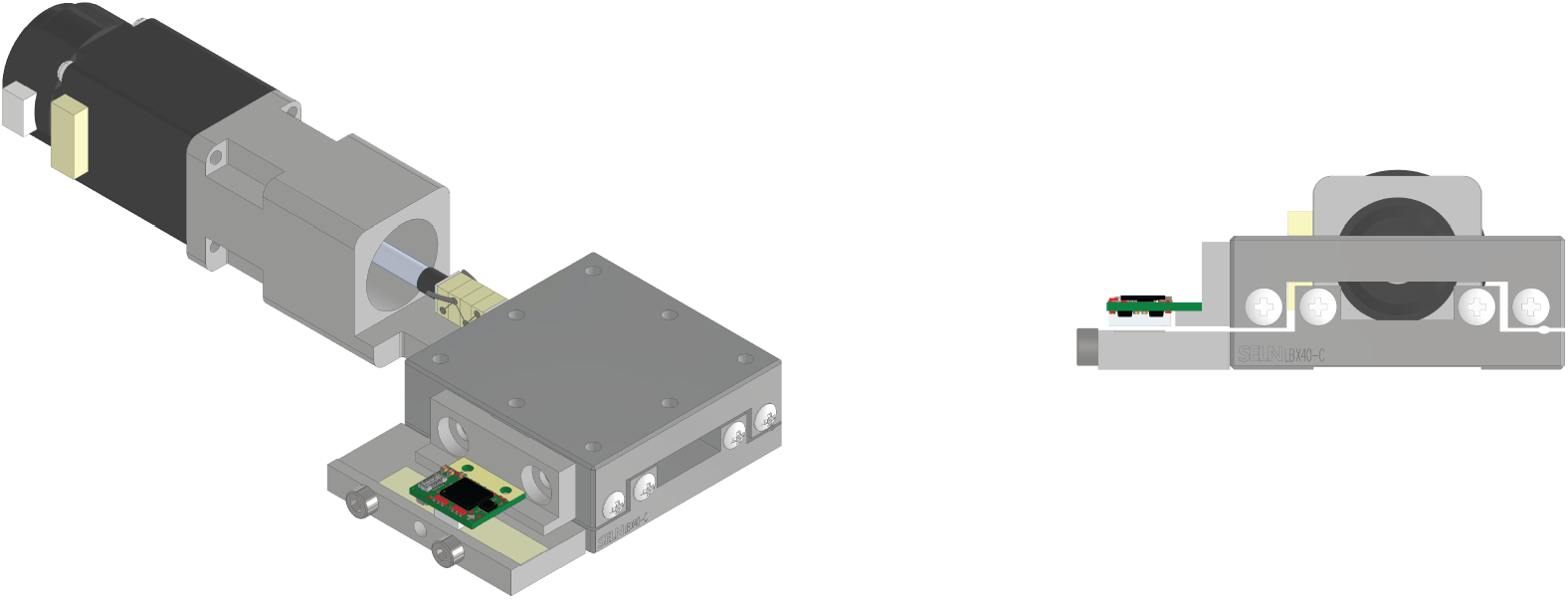
Motorized z-stage with an integrated optical encoder (perspective and front view). The tape and encoder are mounted on CNC machined adapters that are attached to the stage.

**Supplementary Figure 3.**
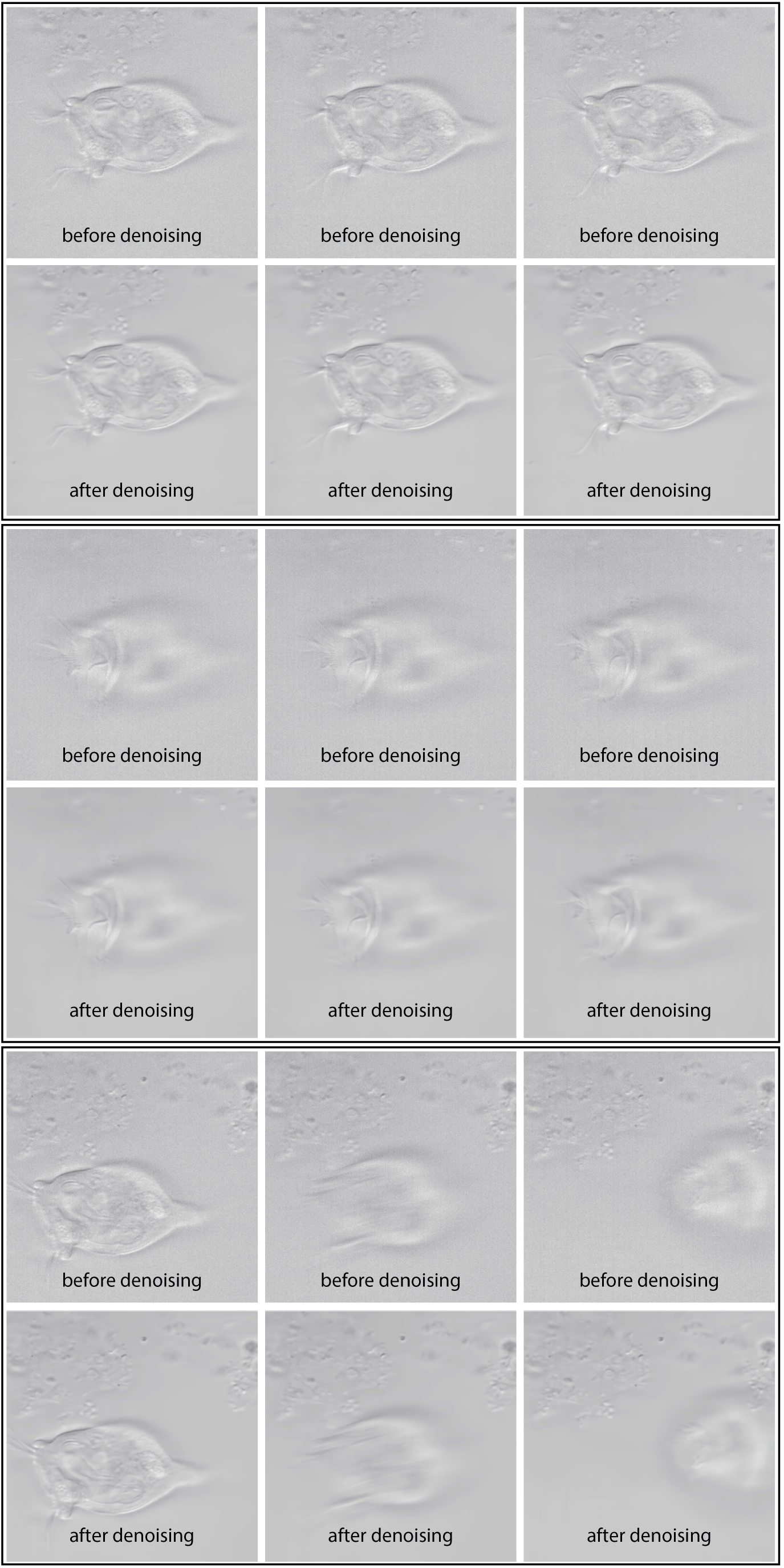
Comparisons of images from Figure 6 (On Semiconductor Python 300 CMOS sensor, LED matrix illumination, 20 *µ*s exposure time) before and after FFDNet denoising. FFDNet denoising effectively reduced noise and at the same time preserved details. We found the same denoising also worked well with images from fluorescence microscopy using color CMOS sensors [44].

**Supplementary Figure 4.**
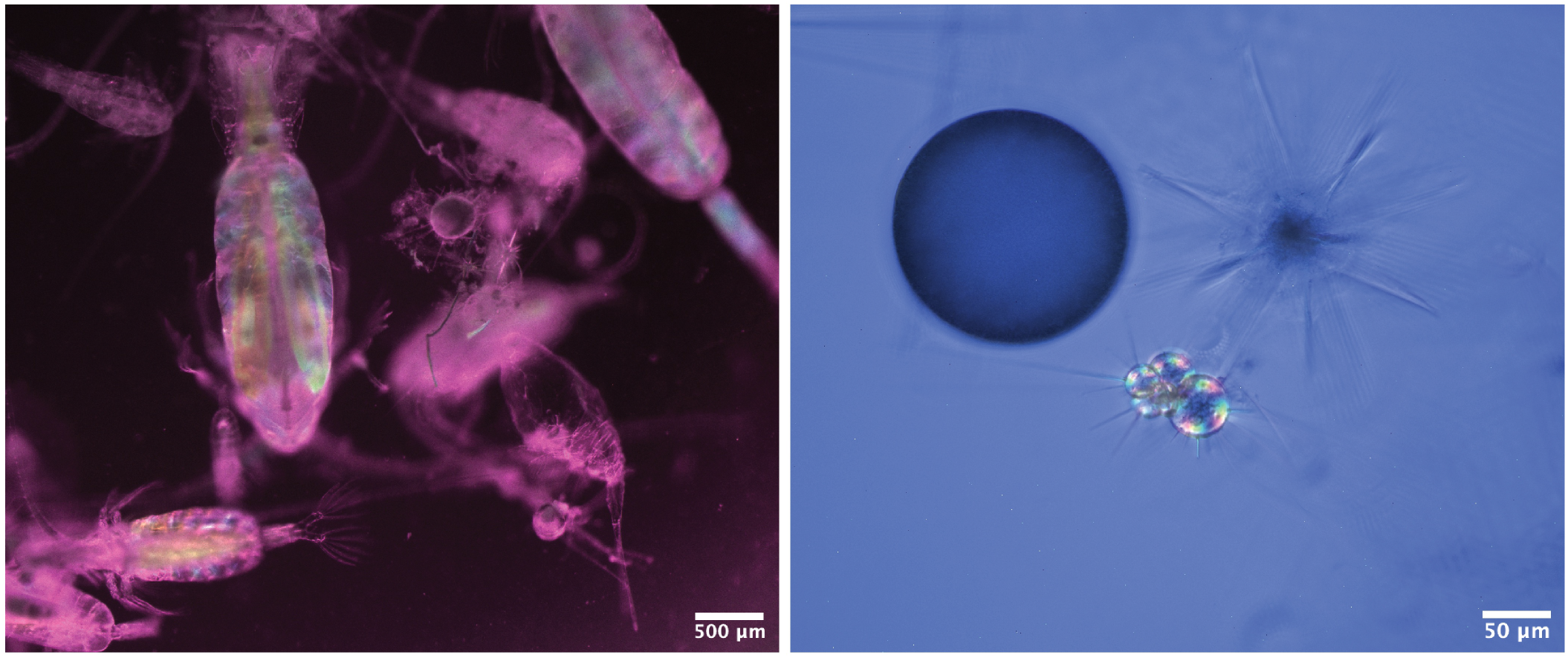
Single-shot polarization imaging with Sony IMX250MZR of Copepod (A) and Forminefera (B). The images were collected while we were on board R/V Kilo Moana during the HOT-317 cruise (Dec 18-22, 2019) (Supplementary Figure 5).

**Supplementary Figure 5.**
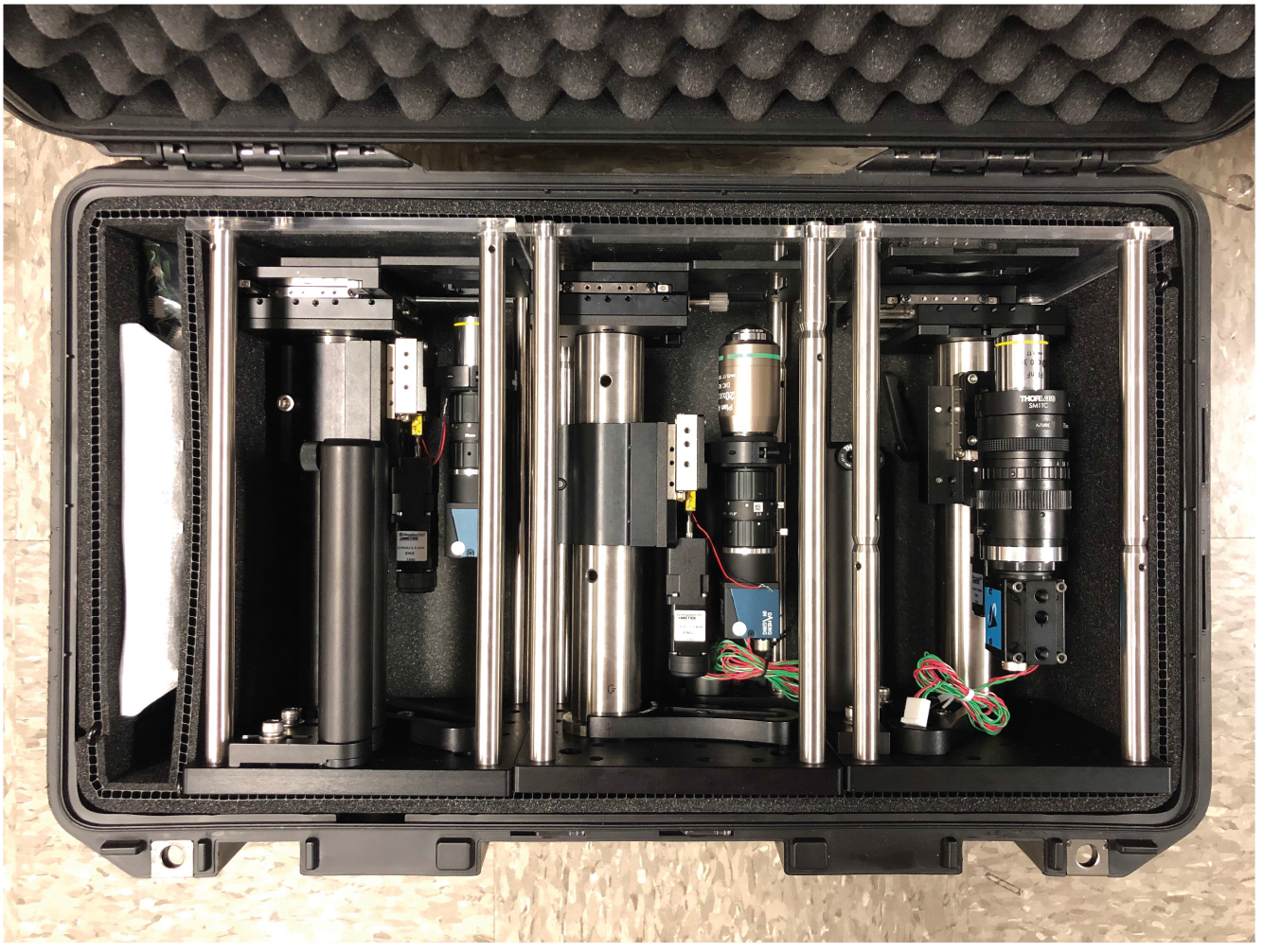
Three Squid microscopes in a Pelican Air 1535 carry-on case.

**Supplementary Table 1:**
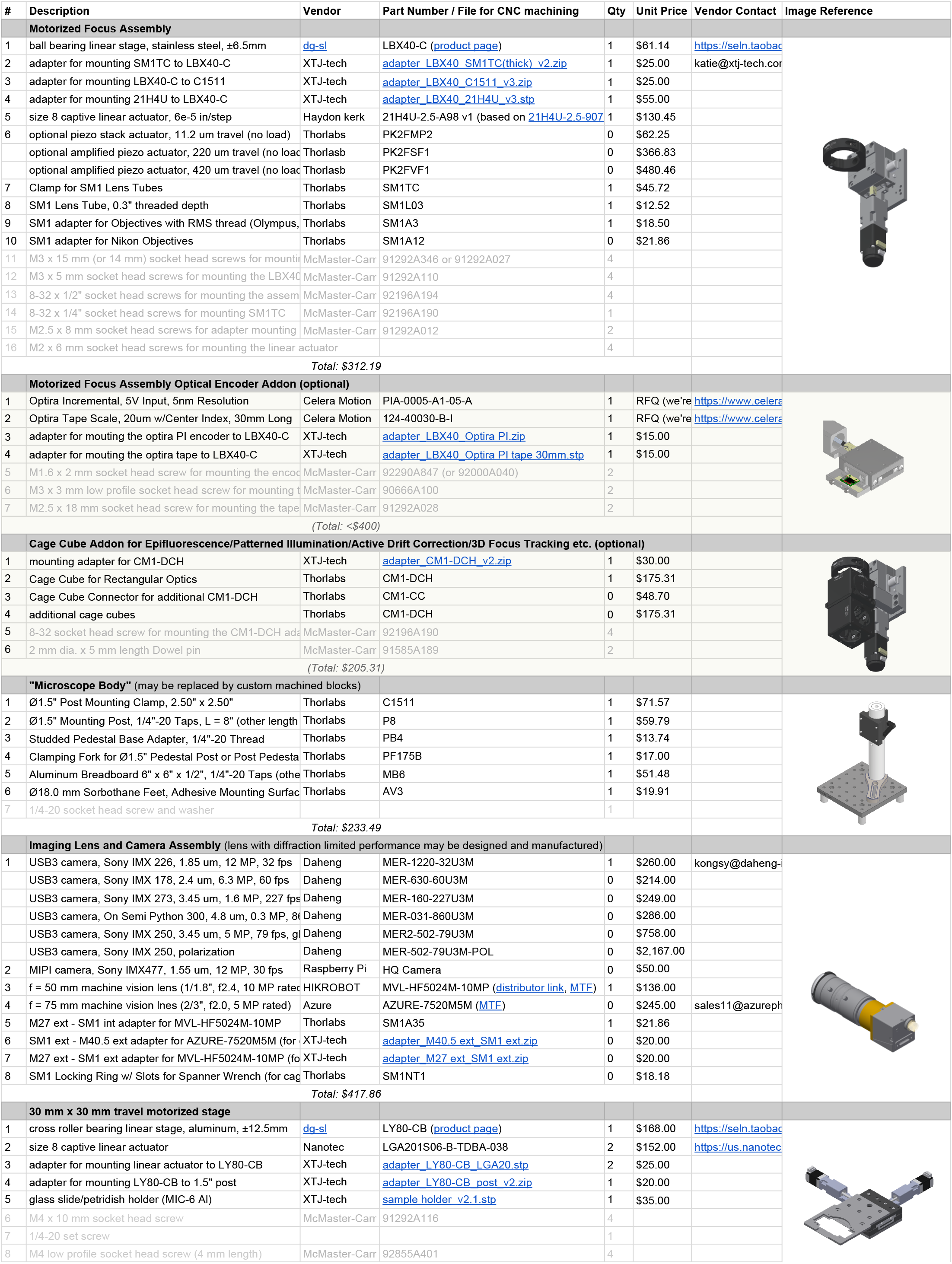
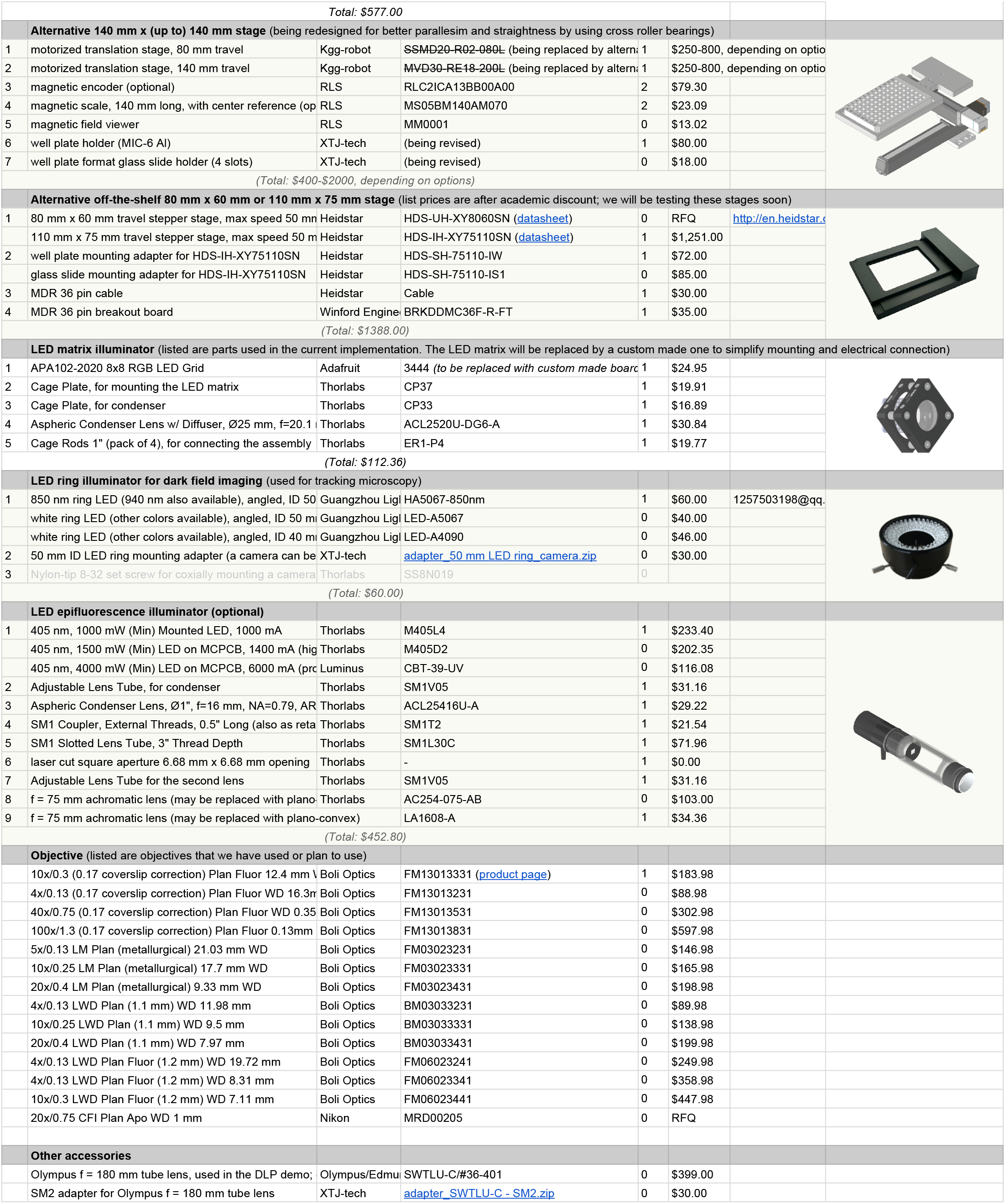
Microscope BOM.

**Supplementary Table 2:**
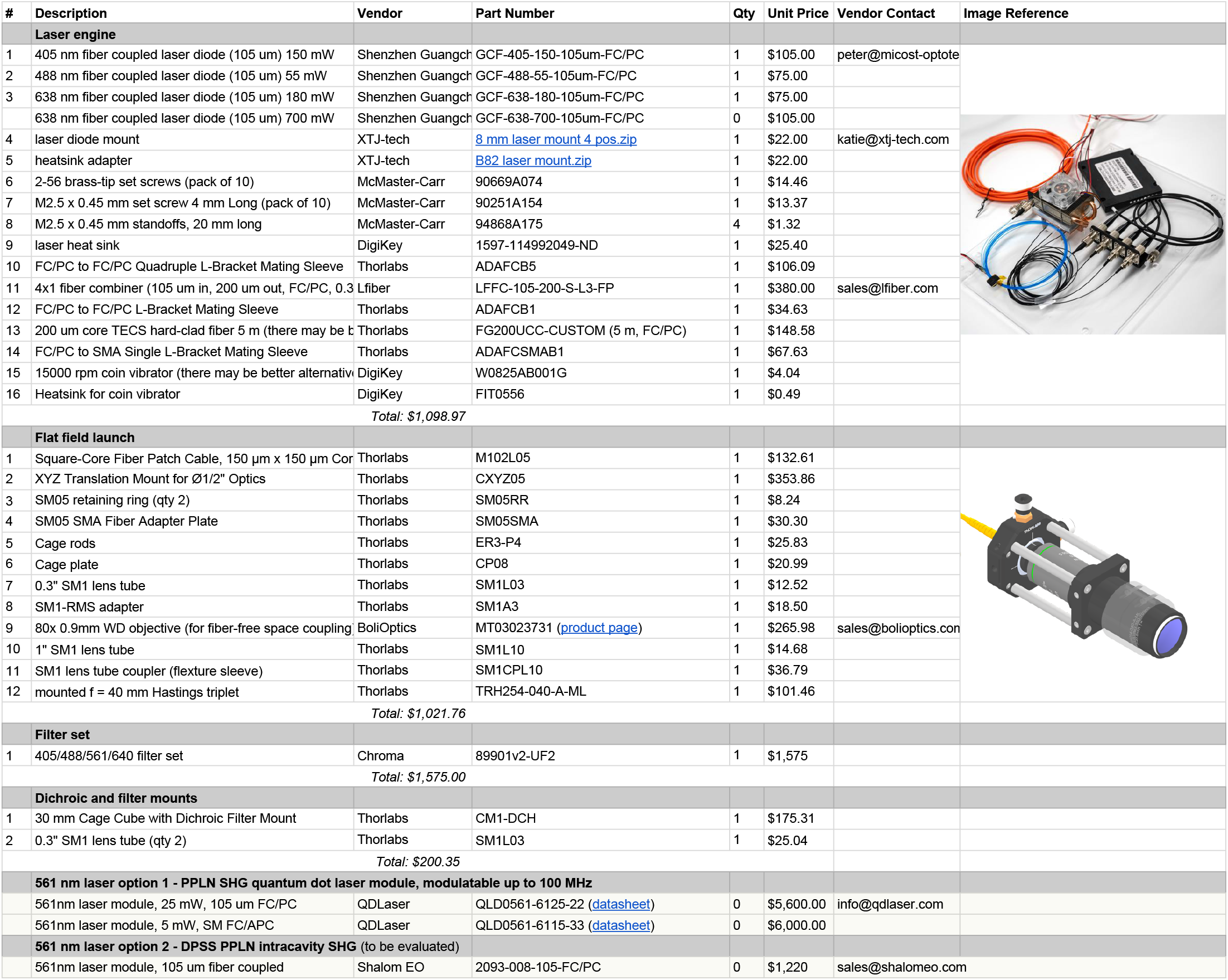
Laser Engine and Flat field launch BOM.

**Supplementary Table 3:**
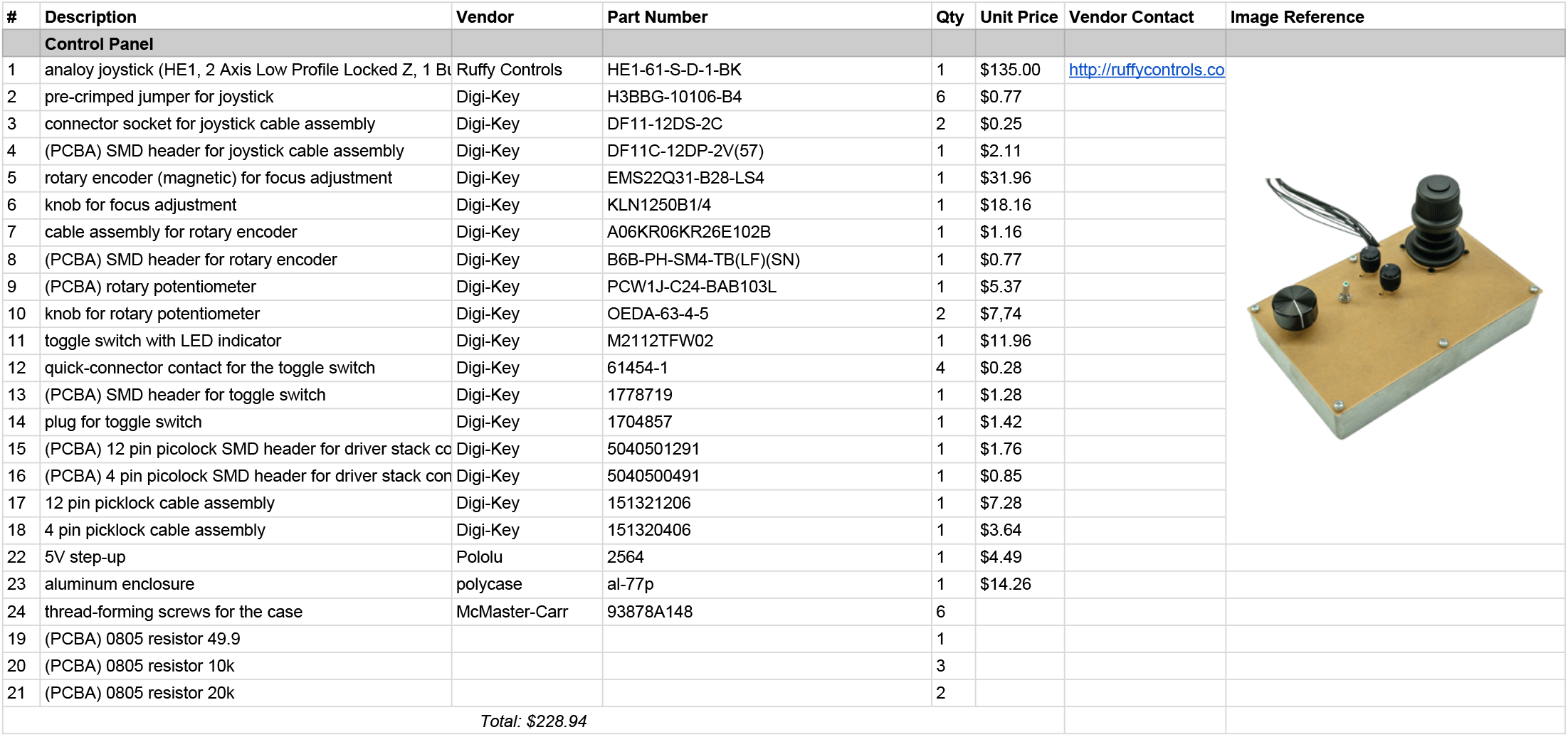
Control Panel BOM.

| # | Description | Vendor | Part Number | Qty | Unit Price | Vendor Contact | Image Reference |
| --- | --- | --- | --- | --- | --- | --- | --- |
| **Control Panel** |  |  |  |  |  |  |  |
| 1 | analog joystick (HE1, 2 Axis Low Profile Locked Z, 1 Button) | Ruffly Controls | HE1-61-S-D-1-BK | 1 | \$135.00 | http://rufflycontrols.com | |
| 2 | pre-crimped jumper for joystick | Digi-Key | H3BBG-10106-B4 | 6 | \$0.77 | | |
| 3 | connector socket for joystick cable assembly | Digi-Key | DF11-12DS-2C | 2 | \$0.25 | | |
| 4 | (PCBA) SMD header for joystick cable assembly | Digi-Key | DF11C-12DP-2V(57) | 1 | \$2.11 | | |
| 5 | rotary encoder (magnetic) for focus adjustment | Digi-Key | EMS22Q31-B28-LS4 | 1 | \$31.96 | | |
| 6 | knob for focus adjustment | Digi-Key | KLN1250B1/4 | 1 | \$18.16 | | |
| 7 | cable assembly for rotary encoder | Digi-Key | A06KR06KR26E102B | 1 | \$1.16 | | |
| 8 | (PCBA) SMD header for rotary encoder | Digi-Key | B6B-PH-SM4-TB(LF)(SN) | 1 | \$0.77 | | |
| 9 | (PCBA) rotary potentiometer | Digi-Key | PCW1J-C24-BAB103L | 1 | \$5.37 | | |
| 10 | knob for rotary potentiometer | Digi-Key | OEDA-63-4-5 | 2 | \$7.74 | | |
| 11 | toggle switch with LED indicator | Digi-Key | M2112TFW02 | 1 | \$11.96 | | |
| 12 | quick-connector contact for the toggle switch | Digi-Key | 61454-1 | 4 | \$0.28 | | |
| 13 | (PCBA) SMD header for toggle switch | Digi-Key | 1778719 | 1 | \$1.28 | | |
| 14 | plug for toggle switch | Digi-Key | 1704857 | 1 | \$1.42 | | |
| 15 | (PCBA) 12 pin picolock SMD header for driver stack control | Digi-Key | 5040501291 | 1 | \$1.76 | | |
| 16 | (PCBA) 4 pin picolock SMD header for driver stack control | Digi-Key | 5040500491 | 1 | \$0.85 | | |
| 17 | 12 pin picklock cable assembly | Digi-Key | 151321206 | 1 | \$7.28 | | |
| 18 | 4 pin picklock cable assembly | Digi-Key | 151320406 | 1 | \$3.64 | | |
| 22 | 5V step-up | Pololu | 2564 | 1 | \$4.49 | | |
| 23 | aluminum enclosure | polycase | al-77p | 1 | \$14.26 | | |
| 24 | thread-forming screws for the case | McMaster-Carr | 93878A148 | 6 |  |  |  |
| 19 | (PCBA) 0805 resistor 49.9 |  |  | 1 |  |  |  |
| 20 | (PCBA) 0805 resistor 10k |  |  | 3 |  |  |  |
| 21 | (PCBA) 0805 resistor 20k |  |  | 2 |  |  |  |
| | | | | | **Total: \$228.94** | | |

## References

1. Mahecic Dora, Gambarotto Davide, M Douglass Kyle, Fortun Denis, Banterle Niccoló, A Ibrahim Khalid, Le Guennec Maeva, Gönczy Pierre, Hamel Virginie, Guichard Paul et al., “Homogeneous Multifocal Excitation for High-Throughput Super-Resolution Imaging,” Nat. methods.

2. Alexander Auer, Thomas Schlichthaerle, Johannes B Woehrstein, Florian Schueder, Maximilian T Strauss, Heinrich Grabmayr, and Ralf Jungmann, “Nanometer-scale Multiplexed Super-Resolution Imaging with an Economic 3D-DNA-PAINT Microscope,” ChemPhysChem 19, 3024–3034 (2018).

3. Jonatan Alvelid and Ilaria Testa, “Stable stimulated emission depletion imaging of extended sample regions,” J. Phys. D: Appl. Phys. 53, 024001 (2019).

4. Anna-Karin Gustavsson, Petar N Petrov, Maurice Y Lee, Yoav Shechtman, and WE Moerner, “3D single-molecule super-resolution microscopy with a tilted light sheet,” Nat. communications 9, 1–8 (2018).

5. Kwasi Kwakwa, Alexander Savell, Timothy Davies, Ian Munro, Simona Parrinello, Marco A Purbhoo, Chris Dunsby, Mark AA Neil, and Paul MW French, “easySTORM: a robust, lower-cost approach to localisation and TIRF microscopy,” J. Biophotonics 9, 948–957 (2016).

6. Luciano A Masullo, Andreas Bodén, Francesca Pennacchietti, Giovanna Coceano, Michael Ratz, and Ilaria Testa, “Enhanced photon collection enables four dimensional fluorescence nanoscopy of living systems,” Nat. communications 9, 1–9 (2018).

7. Andrew ES Barentine, Yu Lin, Miao Liu, Phylicia Kidd, Leonhard Balduf, Michael R Grace, Siyuan Wang, Joerg Bewersdorf, and David Baddeley, “3D Multicolor Nanoscopy at 10,000 Cells a Day,” bioRxiv p. 606954 (2019).

8. Koen JA Martens, Sam PB van Beljouw, Simon van der Els, Jochem NA Vink, Sander Baas, George A Vogelaar, Stan JJ Brouns, Peter van Baarlen, Michiel Kleerebezem, and Johannes Hohlbein, “Visualisation of dCas9 target search in vivo using an open-microscopy framework,” Nat. communications 10, 1–11 (2019).

9. Maja Klevanski, Frank Herrmannsdoerfer, Steffen Sass, Varun Venkataramani, Mike Heilemann, and Thomas Kuner, “Automated highly multiplexed super-resolution imaging of protein nano-architecture in cells and tissues,” Nat. communications 11, 1–11 (2020).

10. Fang Huang, George Sirinakis, Edward S Allgeyer, Lena K Schroeder, Whitney C Duim, Emil B Kromann, Thomy Phan, Felix E Rivera-Molina, Jordan R Myers, Irnov Irnov et al., “Ultra-high resolution 3D imaging of whole cells,” Cell 166, 1028–1040 (2016).

11. Jingyu Wang, Edward S Allgeyer, George Sirinakis, Yongdeng Zhang, Kevin Hu, Mark D Lessard, Yiming Li, Robin Diekmann, Michael A Phillips, Ian M Dobbie et al., “Implementation of a 4Pi-SMS super-resolution microscope,” Nat. Protoc. pp. 1–51 (2020).

12. Pierre Jouchet, Clément Cabriel, Nicolas Bourg, Marion Bardou, Christian Pous, FORT Emmanuel, and Sandrine Lévêque-Fort, “Nanometric axial localization of single fluorescent molecules with modulated excitation,” BioRxiv p. 865865 (2019).

13. Loïc Reymond, Johannes Ziegler, Christian Knapp, Fung-Chen Wang, Thomas Huser, Verena Ruprecht, and Stefan Wieser, “SIMPLE: Structured illumination based point localization estimator with enhanced precision,” Opt. express 27, 24578–24590 (2019).

14. Lusheng Gu, Yuanyuan Li, Shuwen Zhang, Yanhong Xue, Weixing Li, Dong Li, Tao Xu, and Wei Ji, “Molecular resolution imaging by repetitive optical selective exposure,” Nat. methods 16, 1114–1118 (2019).

15. Jelmer Cnossen, Taylor Hinsdale, Rasmus Ø Thorsen Marĳn Siemons, Florian Schueder, Ralf Jungmann, Carlas S Smith, Bernd Rieger, and Sjoerd Stallinga, “Localization microscopy at doubled precision with patterned illumination,” Nat. methods 17, 59–63 (2020).

16. Klaus C Gwo sch, Jasmin K Pape, Francisco Balzarotti, Philipp Hoess, Jan Ellenberg, Jonas Ries, and Stefan W Hell, “MINFLUX nanoscopy delivers 3D multicolor nanometer resolution in cells,” Nat. methods 17, 217–224 (2020).

17. Luciano A Masullo, Florian Steiner, Jonas Zähringer, Lucía F Lopez, Johann Bohlen, Lars Richter, Fiona Cole, Philip Tinnefeld, and Fernando D Stefani, “Pulsed Interleaved MINFLUX,” Nano Lett. (2020).

18. Daniel Schröder, Joran Deschamps, Anindita Dasgupta, Ulf Matti, and Jonas Ries, “Cost-efficient open source laser engine for microscopy,” Biomed. Opt. Express 11, 609–623 (2020).

19. Dan Dan, Ming Lei, Baoli Yao, Wen Wang, Martin Winterhalder, Andreas Zumbusch, Yujiao Qi, Liang Xia, Shaohui Yan, Yanlong Yang et al., “DMD-based LED-illumination super-resolution and optical sectioning microscopy,” Sci. reports 3, 1116 (2013).

20. Andreas Markwirth, Mario Lachetta, Viola Mönkemöller, Rainer Heintzmann, Wolfgang Hübner, Thomas Huser, and Marcel Müller, “Video-rate multi-color structured illumination microscopy with simultaneous real-time reconstruction,” Nat. communications 10, 1–11 (2019).

21. Jakub Pospíšil, Tomáš Lukeš, Justin Bendesky, Karel Fliegel, Kathrin Spendier, and Guy M Hagen, “Imaging tissues and cells beyond the diffraction limit with structured illumination microscopy and Bayesian image reconstruction,” GigaScience 8, giy126 (2019).

22. Karl A Johnson and Guy M Hagen, “Artifact-free whole-slide imaging with structured illumination microscopy and Bayesian image reconstruction,” GigaScience 9, giaa035 (2020).

23. Benjamin Ambrose, James M Baxter, John Cully, Matthew Willmott, Elliot M Steele, Benji C Bateman, Marisa L Martin-Fernandez, Ashley Cadby, Jonathan Shewring, Marleen Aaldering et al., “The smfBox is an open-source platform for single-molecule FRET,” Nat. communications 11, 1–6 (2020).

24. Christopher A Werley, Miao-Ping Chien, and Adam E Cohen, “Ultrawidefield microscope for high-speed fluorescence imaging and targeted optogenetic stimulation,” Biomed. optics express 8, 5794–5813 (2017).

25. Vicente J Parot, Carlos Sing-Long, Yoav Adam, Urs L Böhm, Linlin Z Fan, Samouil L Farhi, and Adam E Cohen, “Compressed hadamard microscopy for high-speed optically sectioned neuronal activity recordings,” J. Phys. D: Appl. Phys. 52, 144001 (2019).

26. Samouil L Farhi, Vicente J Parot, Abhinav Grama, Masahito Yamagata, Ahmed S Abdelfattah, Yoav Adam, Shan Lou, Jeong Jun Kim, Robert E Campbell, David D Cox et al., “Wide-area all-optical neurophysiology in acute brain slices,” J. Neurosci. 39, 4889–4908 (2019).

27. Cuong Nguyen, Hansini Upadhyay, Michael Murphy, Gabriel Borja, Emily J Rozsahegyi, Adam Barnett, Ted Brookings, Owen B McManus, and Christopher A Werley, “Simultaneous voltage and calcium imaging and optogenetic stimulation with high sensitivity and a wide field of view,” Biomed. optics express 10, 789–806 (2019).

28. Qinyi Fu, Benjamin L Martin, David Q Matus, and Liang Gao, “Imaging multicellular specimens with real-time optimized tiling light-sheet selective plane illumination microscopy,” Nat. communications 7, 1–10 (2016).

29. Yanlu Chen, Xiaoliang Li, Dongdong Zhang, Chunhui Wang, Ruili Feng, Xuzhao Li, Yao Wen, Hao Xu, Xinyi Shirley Zhang, Xiao Yang et al., “A Versatile Tiling Light Sheet Microscope for Imaging of Cleared Tissues,” Cell Reports 33, 108349 (2020).

30. Tonmoy Chakraborty, Meghan K Driscoll, Elise Jeffery, Malea M Murphy, Philippe Roudot, Bo-Jui Chang, Saumya Vora, Wen Mai Wong, Cara D Nielson, Hua Zhang et al., “Light-sheet microscopy of cleared tissues with isotropic, subcellular resolution,” Nat. methods 16, 1109–1113 (2019).

31. Fabian F Voigt, Daniel Kirschenbaum, Evgenia Platonova, Stéphane Pagès, Robert AA Campbell, Rahel Kastli, Martina Schaettin, Ladan Egolf, Alexander Van Der Bourg, Philipp Bethge et al., “The mesoSPIM initiative: open-source light-sheet microscopes for imaging cleared tissue,” Nat. methods 16, 1105–1108 (2019).

32. Reto Fiolka Amir Fardad Lachlan Whitehead Alfred Millett-Sikking, Kevin M. Dean and Andrew York, “AndrewGYork/high_na_single_objective_lightsheet: Work-in-progress,” (2019).

33. Bin Yang, Xingye Chen, Yina Wang, Siyu Feng, Veronica Pessino, Nico Stuurman, Nathan H Cho, Karen W Cheng, Samuel J Lord, Linfeng Xu et al., “Epi-illumination SPIM for volumetric imaging with high spatial-temporal resolution,” Nat. methods 16, 501–504 (2019).

34. Adam K Glaser, Nicholas P Reder, Ye Chen, Chengbo Yin, Linpeng Wei, Soyoung Kang, Lindsey A Barner, Weisi Xie, Erin F McCarty, Chenyi Mao et al., “Multi-immersion open-top light-sheet microscope for high-throughput imaging of cleared tissues,” Nat. communications 10, 1–8 (2019).

35. Venkatakaushik Voleti, Kripa B Patel, Wenze Li, Citlali Perez Campos, Srinidhi Bharadwaj, Hang Yu, Caitlin Ford, Malte J Casper, Richard Wenwei Yan, Wenxuan Liang et al., “Real-time volumetric microscopy of in vivo dynamics and large-scale samples with SCAPE 2.0,” Nat. methods 16, 1054–1062 (2019).

36. Bo-Jui Chang, Mark Kittisopikul, Kevin M Dean, Philippe Roudot, Erik S Welf, and Reto Fiolka, “Universal light-sheet generation with field synthesis,” Nat. methods 16, 235–238 (2019).

37. Etai Sapoznik, Bo-Jui Chang, Jaewon Huh, Robert J Ju, Evgenia V Azarova, Theresa Pohlkamp, Erik S Welf, David Broadbent, Alexandre F Carisey, Samantha J Stehbens et al., “A versatile Oblique Plane Microscope for large-scale and high-resolution imaging of subcellular dynamics,” Elife 9, e57681 (2020).

38. Bin Yang, Alfred Millett-Sikking, Merlin Lange, Ahmet Can Solak, Hirofumi Kobayashi, Andrew York, and Loic A Royer, “High-Resolution, Large Field-of-View, and Multi-View Single Objective Light-Sheet Microscopy,” bioRxiv (2020).

39. Bo-Jui Chang, Etai Sapoznik, Theresa Pohlkamp, Tamara Terrones, Erik S Welf, James David Manton, Philippe Roudot, Kayley Hake, Lachlan Whitehead, Andrew G York et al., “Real-Time Multi-Angle Projection Imaging of Biological Dynamics,” bioRxiv (2020).

40. Qingshan Wei, Hangfei Qi, Wei Luo, Derek Tseng, So Jung Ki, Zhe Wan, Zoltán Göröcs, Laurent A Bentolila, Ting-Ting Wu, Ren Sun et al., “Fluorescent imaging of single nanoparticles and viruses on a smart phone,” ACS nano 7, 9147–9155 (2013).

41. James S Cybulski, James Clements, and Manu Prakash, “Foldscope: origami-based paper microscope,” PloS one 9, e98781 (2014).

42. Arunan Skandarajah, Clay D Reber, Neil A Switz, and Daniel A Fletcher, “Quantitative imaging with a mobile phone microscope,” PloS one 9, e96906 (2014).

43. Michael V D’Ambrosio, Matthew Bakalar, Sasisekhar Bennuru, Clay Reber, Arunan Skandarajah, Lina Nilsson, Neil Switz, Joseph Kamgno, Sébastien Pion, Michel Boussinesq et al., “Point-of-care quantification of blood-borne filarial parasites with a mobile phone microscope,” Sci. translational medicine 7, 286re4–286re4 (2015).

44. Hongquan Li, Hazel Soto-Montoya, Maxime Voisin, Lucas Fuentes Valenzuela, and Manu Prakash, “Octopi: Open configurable high-throughput imaging platform for infectious disease diagnosis in the field,” bioRxiv p. 684423 (2019).

45. Tomas Aidukas, Regina Eckert, Andrew R Harvey, Laura Waller, and Pavan C Konda, “Low-cost, sub-micron resolution, wide-field computational microscopy using opensource hardware,” Sci. reports 9, 1–12 (2019).

46. Chengfei Guo, Zichao Bian, Shaowei Jiang, Michael Murphy, Jiakai Zhu, Ruihai Wang, Pengming Song, Xiaopeng Shao, Yongbing Zhang, and Guoan Zheng, “OpenWSI: a low-cost, high-throughput whole slide imaging system via single-frame autofocusing and open-source hardware,” Opt. Lett. 45, 260–263 (2020).

47. Benedict Diederich, René Lachmann, Swen Carlstedt, Barbora Marsikova, Haoran Wang, Xavier Uwurukundo, Alexander S Mosig, and Rainer Heintzmann, “A versatile and customizable low-cost 3D-printed open standard for microscopic imaging,” Nat. Commun. 11, 1–9 (2020).

48. Thibaut Pollina, Adam Larson, Fabien Lombard, Hongquan Li, Sebastien Colin, Colomban de Vargas, and Manu Prakash, “PlanktonScope: Affordable modular imaging platform for citizen oceanography,” bioRxiv (2020).

49. James P Sharkey, Darryl CW Foo, Alexandre Kabla, Jeremy J Baumberg, and Richard W Bowman, “A one-piece 3D printed flexure translation stage for open-source microscopy,” Rev. Sci. Instruments 87, 025104 (2016).

50. Joel T Collins, Joe Knapper, Julian Stirling, Joram Mduda, Catherine Mkindi, Valeriana Mayagaya, Grace A Mwakajinga, Paul T Nyakyi, Valerian L Sanga, Dave Carbery et al., “Robotic microscopy for everyone: the OpenFlexure Microscope,” Biomed. Opt. Express 11, 2447–2460 (2020).

51. Jeffrey P Nguyen, Frederick B Shipley, Ashley N Linder, George S Plummer, Mochi Liu, Sagar U Setru, Joshua W Shaevitz, and Andrew M Leifer, “Whole-brain calcium imaging with cellular resolution in freely behaving Caenorhabditis elegans,” Proc. Natl. Acad. Sci. 113, E1074–E1081 (2016).

52. Dal Hyung Kim, Jungsoo Kim, João C Marques, Abhinav Grama, David GC Hildebrand, Wenchao Gu, Jennifer M Li, and Drew N Robson, “Pan-neuronal calcium imaging with cellular resolution in freely swimming zebrafish,” Nat. methods 14, 1107–1114 (2017).

53. Lin Cong, Zeguan Wang, Yuming Chai, Wei Hang, Chunfeng Shang, Wenbin Yang, Lu Bai, Jiulin Du, Kai Wang, and Quan Wen, “Rapid whole brain imaging of neural activity in freely behaving larval zebrafish (Danio rerio),” Elife 6, e28158 (2017).

54. Deepak Krishnamurthy, Hongquan Li, François Benoit du Rey, Pierre Cambournac, Adam G Larson, Ethan Li, and Manu Prakash, “Scale-free vertical tracking microscopy,” Nat. Methods pp. 1–12 (2020).

55. Guoan Zheng, Roarke Horstmeyer, and Changhuei Yang, “Wide-field, high-resolution Fourier ptychographic microscopy,” Nat. photonics 7, 739–745 (2013).

56. Lei Tian, Zĳi Liu, Li-Hao Yeh, Michael Chen, Jingshan Zhong, and Laura Waller, “Computational illumination for high-speed in vitro Fourier ptychographic microscopy,” Optica 2, 904–911 (2015).

57. Lei Tian and Laura Waller, “Quantitative differential phase contrast imaging in an LED array microscope,” Opt. express 23, 11394–11403 (2015).

58. Michael Chen, Lei Tian, and Laura Waller, “3D differential phase contrast microscopy,” Biomed. optics express 7, 3940–3950 (2016).

59. Yao Fan, Jiasong Sun, Qian Chen, Xiangpeng Pan, Maciej Trusiak, and Chao Zuo, “Single-shot isotropic quantitative phase microscopy based on color-multiplexed differential phase contrast,” APL Photonics 4, 121301 (2019).

60. Michael R Kellman, Emrah Bostan, Nicole A Repina, and Laura Waller, “Physics-based learned design: Optimized coded-illumination for quantitative phase imaging,” IEEE Transactions on Comput. Imaging 5, 344–353 (2019).

61. Jiaji Li, Alex Matlock, Yunzhe Li, Qian Chen, Chao Zuo, and Lei Tian, “High-speed in vitro intensity diffraction tomography,” Adv. Photonics 1, 066004 (2019).

62. Shwetadwip Chowdhury, Michael Chen, Regina Eckert, David Ren, Fan Wu, Nicole Repina, and Laura Waller, “High-resolution 3D refractive index microscopy of multiple-scattering samples from intensity images,” Optica 6, 1211–1219 (2019).

63. Li-Hao Yeh, Shwetadwip Chowdhury, and Laura Waller, “Computational structured illumination for high-content fluorescence and phase microscopy,” Biomed. optics express 10, 1978–1998 (2019).

64. Fanglin Linda Liu, Grace Kuo, Nick Antipa, Kyrollos Yanny, and Laura Waller, “Fourier diffuserScope: single-shot 3D Fourier light field microscopy with a diffuser,” Opt. Express 28, 28969–28986 (2020).

65. Celalettin Yurdakul, Oguzhan Avci, Alex Matlock, Alexander J Devaux, Maritza V Quintero, Ekmel Ozbay, Robert A Davey, John H Connor, W Clem Karl, Lei Tian et al., “High-throughput, high-resolution interferometric light microscopy of biological nanoparticles,” ACS nano 14, 2002–2013 (2020).

66. Syuan-Ming Guo, Li-Hao Yeh, Jenny Folkesson, Ivan E Ivanov, Anitha P Krishnan, Matthew G Keefe, Ezzat Hashemi, David Shin, Bryant B Chhun, Nathan H Cho et al., “Revealing architectural order with quantitative label-free imaging and deep learning,” Elife 9, e55502 (2020).

67. Li-Hao Yeh, Ivan E. Ivanov, Bryant B. Chhun, Syuan-Ming Guo, Ezzat Hashemi, Janie R. Byrum, Juan A. Pérez-Bermejo, Huĳun Wang, Yanhao Yu, Peter G. Kazansky, Bruce R. Conklin, May H. Han, and Shalin B. Mehta, “uPTI: uniaxial permittivity tensor imaging of intrinsic density and anisotropy,” bioRxiv (2020).

68. Shiyi Cheng, Sipei Fu, Yumi Mun Kim, Weiye Song, Yunzhe Li, Yujia Xue, Ji Yi, and Lei Tian, “Single-cell cytometry via multiplexed fluorescence prediction by label-free reflectance microscopy,” bioRxiv (2020).

69. Chao Zuo, Jiasong Sun, Jiaji Li, Anand Asundi, and Qian Chen, “Wide-field high-resolution 3d microscopy with fourier ptychographic diffraction tomography,” Opt. Lasers Eng. 128, 106003 (2020).

70. Lingbo Jin, Yubo Tang, Yicheng Wu, Jackson B Coole, Melody T Tan, Xuan Zhao, Hawraa Badaoui, Jacob T Robinson, Michelle D Williams, Ann M Gillenwater et al., “Deep learning extended depth-of-field microscope for fast and slide-free histology,” Proc. Natl. Acad. Sci. (2020).

71. Marcel Müller, Viola Mönkemöller, Simon Hennig, Wolfgang Hübner, and Thomas Huser, “Open-source image reconstruction of super-resolution structured illumination microscopy data in ImageJ,” Nat. communications 7, 1–6 (2016).

72. Artur Speiser, Lucas-Raphael Müller, Ulf Matti, Christopher J Obara, Wesley R Legant, Anna Kreshuk, Jakob H Macke, Jonas Ries, and Srinivas C Turaga, “Deep learning enables fast and dense single-molecule localization with high accuracy,” bioRxiv (2020).

73. Elias Nehme, Daniel Freedman, Racheli Gordon, Boris Ferdman, Lucien E Weiss, Onit Alalouf, Tal Naor, Reut Orange, Tomer Michaeli, and Yoav Shechtman, “DeepSTORM3D: dense 3D localization microscopy and PSF design by deep learning,” Nat. Methods 17, 734–740 (2020).

74. Leonhard Möckl, Anish R Roy, Petar N Petrov, and WE Moerner, “Accurate and rapid background estimation in single-molecule localization microscopy using the deep neural network BGnet,” Proc. Natl. Acad. Sci. 117, 60–67 (2020).

75. Nils Gustafsson, Siân Culley, George Ashdown, Dylan M Owen, Pedro Matos Pereira, and Ricardo Henriques, “Fast live-cell conventional fluorophore nanoscopy with ImageJ through super-resolution radial fluctuations,” Nat. communications 7, 1–9 (2016).

76. Romain F Laine, Kalina L Tosheva, Nils Gustafsson, Robert DM Gray, Pedro Almada, David Albrecht, Gabriel T Risa, Fredrik Hurtig, Ann-Christin Lindås, Buzz Baum et al., “NanoJ: a high-performance open-source super-resolution microscopy toolbox,” J. Phys. D: Appl. Phys. 52, 163001 (2019).

77. Min Guo, Yue Li, Yĳun Su, Talley Lambert, Damian Dalle Nogare, Mark W Moyle, Leighton H Duncan, Richard Ikegami, Anthony Santella, Ivan Rey-Suarez et al., “Rapid image deconvolution and multiview fusion for optical microscopy,” Nat. Biotechnol. pp. 1–10 (2020).

78. Martin Weigert, Uwe Schmidt, Tobias Boothe, Andreas Müller, Alexandr Dibrov, Akanksha Jain, Benjamin Wilhelm, Deborah Schmidt, Coleman Broaddus, Siân Culley et al., “Content-aware image restoration: pushing the limits of fluorescence microscopy,” Nat. methods 15, 1090–1097 (2018).

79. Linjing Fang, Fred Monroe, Sammy Weiser Novak, Lyndsey Kirk, Cara R Schiavon, B Yu Seungyoon, Tong Zhang, Melissa Wu, Kyle Kastner, Yoshiyuki Kubota et al., “Deep learning-based point-scanning super-resolution imaging,” bioRxiv p. 740548 (2019).

80. Hongda Wang, Yair Rivenson, Yiyin Jin, Zhensong Wei, Ronald Gao, Harun Günaydin, Laurent A Bentolila, Comert Kural, and Aydogan Ozcan, “Deep learning enables cross-modality super-resolution in fluorescence microscopy,” Nat. methods 16, 103–110 (2019).

81. Yair Rivenson, Hongda Wang, Zhensong Wei, Kevin de Haan, Yibo Zhang, Yichen Wu, Harun Günaydin, Jonathan E Zuckerman, Thomas Chong, Anthony E Sisk et al., “Virtual histological staining of unlabelled tissue-autofluorescence images via deep learning,” Nat. biomedical engineering 3, 466 (2019).

82. Yair Rivenson, Tairan Liu, Zhensong Wei, Yibo Zhang, Kevin de Haan, and Aydogan Ozcan, “PhaseStain: the digital staining of label-free quantitative phase microscopy images using deep learning,” Light. Sci. & Appl. 8, 1–11 (2019).

83. Lucas Von Chamier, Johanna Jukkala, Christoph Spahn, Martina Lerche, Sara Hernández-Pérez, Pieta Mattila, Eleni Karinou, Seamus Holden, Ahmet Can Solak, Alexander Krull et al., “ZeroCostDL4Mic: an open platform to simplify access and use of Deep-Learning in Microscopy,” BioRxiv (2020).

84. Mark-Anthony Bray, Shantanu Singh, Han Han, Chadwick T Davis, Blake Borgeson, Cathy Hartland, Maria Kost-Alimova, Sigrun M Gustafsdottir, Christopher C Gibson, and Anne E Carpenter, “Cell Painting, a high-content image-based assay for morphological profiling using multiplexed fluorescent dyes,” Nat. protocols 11, 1757 (2016).

85. Claire McQuin, Allen Goodman, Vasiliy Chernyshev, Lee Kamentsky, Beth A Cimini, Kyle W Karhohs, Minh Doan, Liya Ding, Susanne M Rafelski, Derek Thirstrup et al., “CellProfiler 3.0: Next-generation image processing for biology,” PLoS biology 16, e2005970 (2018).

86. Robert Haase, Loic A Royer, Peter Steinbach, Deborah Schmidt, Alexandr Dibrov, Uwe Schmidt, Martin Weigert, Nicola Maghelli, Pavel Tomancak, Florian Jug et al., “CLĲ: GPU-accelerated image processing for everyone,” Nat. Methods 17, 5–6 (2020).

87. Wei Ouyang, Florian Mueller, Martin Hjelmare, Emma Lundberg, and Christophe Zimmer, “ImJoy: an open-source computational platform for the deep learning era,” Nat. methods 16, 1199–1200 (2019).

88. Biagio Mandracchia, Xuanwen Hua, Changliang Guo, Jeonghwan Son, Tara Urner, and Shu Jia, “Fast and accurate sCMOS noise correction for fluorescence microscopy,” Nat. communications 11, 1–12 (2020).

89. Alexander Krull, Tim-Oliver Buchholz, and Florian Jug, “Noise2void-learning denoising from single noisy images,” in Proceedings of the IEEE Conference on Computer Vision and Pattern Recognition, (2019), pp. 2129–2137.

90. Alexander Krull, Tomas Vicar, and Florian Jug, “Probabilistic noise2void: Unsupervised content-aware denoising,” arXiv preprint arXiv:1906.00651 (2019).

91. Mangal Prakash, Manan Lalit, Pavel Tomancak, Alexander Krul, and Florian Jug, “Fully unsupervised probabilistic noise2void,” in 2020 IEEE 17th International Symposium on Biomedical Imaging (ISBI), (IEEE, 2020), pp. 154–158.

92. Tim-Oliver Buchholz, Mangal Prakash, Alexander Krull, and Florian Jug, “DenoiSeg: Joint Denoising and Segmentation,” arXiv preprint arXiv:2005.02987 (2020).

93. Anna S Goncharova, Alf Honigmann, Florian Jug, and Alexander Krull, “Improving Blind Spot Denoising for Microscopy,” arXiv preprint arXiv:2008.08414 (2020).

94. Mangal Prakash, Alexander Krull, and Florian Jug, “DivNoising: Diversity Denoising with Fully Convolutional Variational Autoencoders,” arXiv preprint arXiv:2006.06072 (2020).

95. Uwe Schmidt, Martin Weigert, Coleman Broaddus, and Gene Myers, “Cell Detection with Star-Convex Polygons,” in Medical Image Computing and Computer Assisted Intervention - MICCAI 2018 - 21st International Conference, Granada, Spain, September 16-20, 2018, Proceedings, Part II, (2018), pp. 265–273.

96. Martin Weigert, Uwe Schmidt, Robert Haase, Ko Sugawara, and Gene Myers, “Star-convex Polyhedra for 3D Object Detection and Segmentation in Microscopy,” in The IEEE Winter Conference on Applications of Computer Vision (WACV), (2020).

97. Carsen Stringer, Tim Wang, Michalis Michaelos, and Marius Pachitariu, “Cellpose: a generalist algorithm for cellular segmentation,” Nat. Methods pp. 1–7 (2020).

98. Fabian Isensee, Paul F Jaeger, Simon AA Kohl, Jens Petersen, and Klaus H Maier-Hein, “nnU-Net: a self-configuring method for deep learning-based biomedical image segmentation,” Nat. Methods pp. 1–9 (2020).

99. Zhenqin Wu, Bryant B Chhun, Galina Schmunk, Chang Kim, Li-Hao Yeh, Tomasz J Nowakowski, James Zou, and Shalin B Mehta, “DynaMorph: learning morphodynamic states of human cells with live imaging and sc-RNAseq,” bioRxiv (2020).

100. Assaf Zaritsky, Andrew R Jamieson, Erik S Welf, Andres Nevarez, Justin Cillay, Ugur Eskiocak, Brandi L Cantarel, and Gaudenz Danuser, “Interpretable deep learning of label-free live cell images uncovers functional hallmarks of highly-metastatic melanoma,” BioRxiv (2020).

101. Thomas Blanc, Mohamed El Beheiry, Clément Caporal, Jean-Baptiste Masson, and Bassam Hajj, “Genuage: visualize and analyze multidimensional single-molecule point cloud data in virtual reality,” Nat. Methods pp. 1–3 (2020).

102. Kitching Alexandre Esteban-Ferrer Daniel Handa Anoushka Carr Alexander R. Needham Lisa-Maria Ponjavic Aleks Santos Ana Mafalda McColl James Leterrier Christophe Davis Simon J. Henriques Ricardo Lee Steven F. Spark, Alexander, “vLUME: 3D virtual reality for single-molecule localization microscopy,” Nat. Methods (2020).

103. Nicholas Sofroniew, Talley Lambert, Kira Evans, Juan Nunez-Iglesias, Kevin Yamauchi, Ahmet Can Solak, Grzegorz Bokota, ziyangczi, Genevieve Buckley, Philip Winston, Tony Tung, Draga Doncila Pop, Hector, Jeremy Freeman, Matthias Bussonnier, Peter Boone, Loic Royer, Hagai Har-Gil, Shannon Axelrod, Ariel Rokem, Bryant, Justin Kiggins, Mars Huang, Pranathi Vemuri, Reece Dunham, Trevor Manz, jakirkham, Chris Wood, Alexandre de Siqueira, and Bhavya Chopra, “napari/napari: 0.3.8rc2,” (2020).

104. Philippe Roudot, Wesley R Legant, Qiongjing Zou, Kevin M Dean, Erik S Welf, Ana F David, Daniel W Gerlich, Reto Paul Fiolka, Eric Betzig, and Gaudenz Danuser, “u-track 3D: measuring and interrogating intracellular dynamics in three dimensions.” bioRxiv (2020).

105. Fei Chen, Paul W Tillberg, and Edward S Boyden, “Expansion microscopy,” Science 347, 543–548 (2015).

106. Paul W Tillberg, Fei Chen, Kiryl D Piatkevich, Yongxin Zhao, Chih-Chieh Jay Yu, Brian P English, Linyi Gao, Anthony Martorell, Ho-Jun Suk, Fumiaki Yoshida et al., “Protein-retention expansion microscopy of cells and tissues labeled using standard fluorescent proteins and antibodies,” Nat. biotechnology 34, 987–992 (2016).

107. Rongqin Ke, Marco Mignardi, Alexandra Pacureanu, Jessica Svedlund, Johan Botling, Carolina Wählby, and Mats Nilsson, “In situ sequencing for RNA analysis in preserved tissue and cells,” Nat. methods 10, 857–860 (2013).

108. Je Hyuk Lee, Evan R Daugharthy, Jonathan Scheiman, Reza Kalhor, Joyce L Yang, Thomas C Ferrante, Richard Terry, Sauveur SF Jeanty, Chao Li, Ryoji Amamoto et al., “Highly multiplexed subcellular RNA sequencing in situ,” Science 343, 1360–1363 (2014).

109. Kok Hao Chen, Alistair N Boettiger, Jeffrey R Moffitt, Siyuan Wang, and Xiaowei Zhuang, “Spatially resolved, highly multiplexed RNA profiling in single cells,” Science 348, aaa6090 (2015).

110. Simone Codeluppi, Lars E Borm, Amit Zeisel, Gioele La Manno, Josina A van Lunteren, Camilla I Svensson, and Sten Linnarsson, “Spatial organization of the somatosensory cortex revealed by osmFISH,” Nat. methods 15, 932–935 (2018).

111. Xiao Wang, William E Allen, Matthew A Wright, Emily L Sylwestrak, Nikolay Samusik, Sam Vesuna, Kathryn Evans, Cindy Liu, Charu Ramakrishnan, Jia Liu et al., “Three-dimensional intact-tissue sequencing of single-cell transcriptional states,” Science 361, eaat5691 (2018).

112. Chee-Huat Linus Eng, Michael Lawson, Qian Zhu, Ruben Dries, Noushin Koulena, Yodai Takei, Jina Yun, Christopher Cronin, Christoph Karp, Guo-Cheng Yuan et al., “Transcriptome-scale super-resolved imaging in tissues by RNA seqFISH+,” Nature 568, 235–239 (2019).

113. Huy Q Nguyen, Shyamtanu Chattoraj, David Castillo, Son C Nguyen, Guy Nir, Antonios Lioutas, Elliot A Hershberg, Nuno MC Martins, Paul L Reginato, Mohammed Hannan et al., “3D mapping and accelerated super-resolution imaging of the human genome using in situ sequencing,” Nat. Methods 17, 822–832 (2020).

114. Jia-Ren Lin, Mohammad Fallahi-Sichani, and Peter K Sorger, “Highly multiplexed imaging of single cells using a high-throughput cyclic immunofluorescence method,” Nat. communications 6, 1–7 (2015).

115. Jia-Ren Lin, Benjamin Izar, Shu Wang, Clarence Yapp, Shaolin Mei, Parin M Shah, Sandro Santagata, and Peter K Sorger, “Highly multiplexed immunofluorescence imaging of human tissues and tumors using t-CyCIF and conventional optical microscopes,” Elife 7 (2018).

116. Yury Goltsev, Nikolay Samusik, Julia Kennedy-Darling, Salil Bhate, Matthew Hale, Gustavo Vazquez, Sarah Black, and Garry P Nolan, “Deep profiling of mouse splenic architecture with CODEX multiplexed imaging,” Cell 174, 968–981 (2018).

117. Syuan-Ming Guo, Remi Veneziano, Simon Gordonov, Li Li, Eric Danielson, Karen Perez de Arce, Demian Park, Anthony B Kulesa, Eike-Christian Wamhoff, Paul C Blainey et al., “Multiplexed and high-throughput neuronal fluorescence imaging with diffusible probes,” Nat. communications 10, 1–14 (2019).

118. Sinem K Saka, Yu Wang, Jocelyn Y Kishi, Allen Zhu, Yitian Zeng, Wenxin Xie, Koray Kirli, Clarence Yapp, Marcelo Cicconet, Brian J Beliveau et al., “Immuno-SABER enables highly multiplexed and amplified protein imaging in tissues,” Nat. biotechnology 37, 1080–1090 (2019).

119. Owen Janson Syuan-Ming Guo Jenny Folkesson Bryant B. Chhun Joanna Vinden Ivan E. Ivanov Marcus L. Forst Hongquan Li Adam G. Larson Wesley Wu1 Cristina M. Tato Krista M. McCutcheon Michael J. Peluso Timothy J. Henrich Steven G. Deeks Manu Prakash Bryan Greenhouse John E. Pak Shalin B. Mehta Janie R. Byrum, Eric Waltari, “multiSero: Open multiplex-ELISA platform for analyzing antibody responses to SARS-CoV-2 infection,” bioRxiv (2020).

120. Amy Courtney, Luke M Alvey, George OT Merces, Niamh Burke, and Mark Pickering, “The Flexiscope: a low cost, flexible, convertible and modular microscope with automated scanning and micromanipulation,” Royal Soc. open science 7, 191949 (2018).

121. Rusty Nicovich and Filipe Carvalho, “https://github.com/AllenInstitute/octoDAC,”.

122. Karl Bellve, Clive Standley, Lawrence Lifshitz, and Kevin Fogarty, “Design and implementation of 3D focus stabilization for fluorescence microscopy,” Biophys. J. 106, 606a (2014).

123. Hongqiang Ma and Yang Liu, “Super-resolution localization microscopy: Toward high throughput, high quality, and low cost,” APL Photonics 5, 060902 (2020).

124. Bassam Hajj, Jan Wisniewski, Mohamed El Beheiry, Jĳi Chen, Andrey Revyakin, Carl Wu, and Maxime Dahan, “Whole-cell, multicolor superresolution imaging using volumetric multifocus microscopy,” Proc. Natl. Acad. Sci. 111, 17480–17485 (2014).

125. Hongqiang Ma, Rao Fu, Jianquan Xu, and Yang Liu, “A simple and cost-effective setup for super-resolution localization microscopy,” Sci. reports 7, 1–9 (2017).

126. Simao Coelho, Jongho Baek, Matthew S Graus, James M Halstead, Philip R Nicovich, Kristen Feher, Hetvi Gandhi, J Justin Gooding, and Katharina Gaus, “Ultraprecise single-molecule localization microscopy enables in situ distance measurements in intact cells,” Sci. Adv. 6, eaay8271 (2020).

127. Simao Coelho, Jongho Baek, James Walsh, J Justin Gooding, and Katharina Gaus, “3D active stabilization for single-molecule imaging,” Nat. Protoc. pp. 1–19 (2020).

128. Mariano Bossi, Jonas Fölling, Vladimir N Belov, Vadim P Boyarskiy, Rebecca Medda, Alexander Egner, Christian Eggeling, Andreas Schönle, and Stefan W Hell, “Multicolor far-field fluorescence nanoscopy through isolated detection of distinct molecular species,” Nano letters 8, 2463–2468 (2008).

129. Ilaria Testa, Christian A Wurm, Rebecca Medda, Ellen Rothermel, Claas Von Middendorf, Jonas Fölling, Stefan Jakobs, Andreas Schönle, Stefan W Hell, and Christian Eggeling, “Multicolor fluorescence nanoscopy in fixed and living cells by exciting conventional fluorophores with a single wavelength,” Biophys. journal 99, 2686–2694 (2010).

130. Yongdeng Zhang, Lena K Schroeder, Mark D Lessard, Phylicia Kidd, Jeeyun Chung, Yuanbin Song, Lorena Benedetti, Yiming Li, Jonas Ries, Jonathan B Grimm et al., “Nanoscale subcellular architecture revealed by multicolor three-dimensional salvaged fluorescence imaging,” Nat. methods 17, 225–231 (2020).

131. Kristin S Grußmayer, Stefan Geissbuehler, Adrien Descloux, Tomas Lukes, Marcel Leutenegger, Aleksandra Radenovic, and Theo Lasser, “Spectral cross-cumulants for multicolor super-resolved SOFI imaging,” Nat. communications 11, 1–8 (2020).

132. Sara Abrahamsson, Jĳi Chen, Bassam Hajj, Sjoerd Stallinga, Alexander Y Katsov, Jan Wisniewski, Gaku Mizuguchi, Pierre Soule, Florian Mueller, Claire Dugast Darzacq et al., “Fast multicolor 3D imaging using aberration-corrected multifocus microscopy,” Nat. methods 10, 60–63 (2013).

133. Sara Abrahamsson, Rob Ilic, Jan Wisniewski, Brian Mehl, Liya Yu, Lei Chen, Marcelo Davanco, Laura Oudjedi, Jean-Bernard Fiche, Bassam Hajj et al., “Multifocus microscopy with precise color multi-phase diffractive optics applied in functional neuronal imaging,” Biomed. optics express 7, 855–869 (2016).

134. Sheng Xiao, Howard Gritton, Hua-an Tseng, Dana Zemel, Xue Han, and Jerome Mertz, “High-contrast multifocus microscopy with a single camera and z-splitter prism,” Optica 7, 1477–1486 (2020).

135. Hazen P Babcock, “Multiplane and spectrally-resolved single molecule localization microscopy with industrial grade CMOS cameras,” Sci. reports 8, 1–8 (2018).

136. Adrien Descloux, K. Grußmayer, E Bostan, T Lukes, Arno Bouwens, A Sharipov, S Geissbuehler, A-L Mahul-Mellier, HA Lashuel, M Leutenegger et al., “Combined multi-plane phase retrieval and super-resolution optical fluctuation imaging for 4D cell microscopy,” Nat. Photonics 12, 165–172 (2018).

137. Karl A Johnson, Daniel Noble, Rosa Machado, and Guy M Hagen, “Flexible multiplane structured illumination microscope with a four-camera detector,” bioRxiv (2020).

138. Eduardo Hirata-Miyasaki, Gustav M Pettersson, Khant Zaw, Demis D John, Brian Thibeault, Brandon Lynch, Juliana Hernandez, and Sara Abrahamsson, “Camera-Array 25-Plane Multifocus Microscope For Ultrafast Live 3d Imaging,” in CLEO: Applications and Technology, (Optical Society of America, 2020), pp. JW3P–4.

139. Soheil Mojiri, Sebastian Isbaner, Steffen Mühle, Hongje Jang, Albert J Bae, Ingo Gregor, Azam Gholami, and Joerg Enderlein, “Three-dimensional beating dynamics of chlamydomonas flagella,” bioRxiv (2020).

140. Federico M Barabas, Luciano A Masullo, and Fernando D Stefani, “Note: Tormenta: An open source Python-powered control software for camera based optical microscopy,” Rev. Sci. Instruments 87, 126103 (2016).

141. Alfred Millett-Sikking, Nathaniel H. Thayer, Adam Bohnert, and Andrew G. York, “calico/remote_refocus: Pre-print,” (2018).

142. Robin Diekmann, Katharina Till, Marcel Müller, Matthias Simonis, Mark Schüttpelz, and Thomas Huser, “Characterization of an industry-grade CMOS camera well suited for single molecule localization microscopy–high performance super-resolution at low cost,” Sci. reports 7, 1–10 (2017).

143. Robin Van den Eynde, Alice Sandmeyer, Wim Vandenberg, Sam Duwé, Wolfgang Hübner, Thomas Huser, Peter Dedecker, and Marcel Müller, “Quantitative comparison of camera technologies for cost-effective super-resolution optical fluctuation imaging (SOFI),” J. Physics: Photonics 1, 044001 (2019).

144. Pia Otto, Stephan Bergmann, Alice Sandmeyer, Maxim Dirksen, Oliver Wrede, Thomas Hellweg, and Thomas Huser, “Resolving the internal morphology of core–shell microgels with super-resolution fluorescence microscopy,” Nanoscale Adv. 2, 323–331 (2020).

145. R Oldenbourg and G Mei, “New polarized light microscope with precision universal compensator,” J. microscopy 180, 140–147 (1995).

146. Maki Koike-Tani, Takashi Tominaga, Rudolf Oldenbourg, and Tomomi Tani, “Birefringence changes of dendrites in mouse hippocampal slices revealed with polarizing microscopy,” Biophys. J. (2020).

147. MA Sleigh and D Barlow, “Collection of food by Vorticella,” Transactions Am. Microsc. Soc. pp. 482–486 (1976).

148. MA Sleigh, “The form of beat in cilia of Stentor and Opalina,” J. Exp. Biol. 37, 1–10 (1960).

149. Kai Zhang, Wangmeng Zuo, and Lei Zhang, “FFDNet: Toward a fast and flexible solution for CNN-based image denoising,” IEEE Transactions on Image Process. 27, 4608–4622 (2018).

150. Jun Liao, Yutong Jiang, Zichao Bian, Bahareh Mahrou, Aparna Nambiar, Alexander W Magsam, Kaikai Guo, Shiyao Wang, Yong ku Cho, and Guoan Zheng, “Rapid focus map surveying for whole slide imaging with continuous sample motion,” Opt. letters 42, 3379–3382 (2017).

151. Zachary F Phillips, Regina Eckert, and Laura Waller, “Quasi-dome: A self-calibrated high-na led illuminator for fourier ptychography,” in Imaging Systems and Applications, (Optical Society of America, 2017), pp. IW4E–5.

152. Regina Eckert, Zachary F Phillips, and Laura Waller, “Efficient illumination angle self-calibration in Fourier ptychography,” Appl. optics 57, 5434–5442 (2018).

153. Zeyu Zhao, Bo Xin, Luchang Li, and Zhen-Li Huang, “High-power homogeneous illumination for super-resolution localization microscopy with large field-of-view,” Opt. Express 25, 13382–13395 (2017).

154. Joran Deschamps, Andreas Rowald, and Jonas Ries, “Efficient homogeneous illumination and optical sectioning for quantitative single-molecule localization microscopy,” Opt. express 24, 28080–28090 (2016).

155. Makio Tokunaga, Naoko Imamoto, and Kumiko Sakata-Sogawa, “Highly inclined thin illumination enables clear single-molecule imaging in cells,” Nat. methods 5, 159–161 (2008).

156. Daryl Lim, Kengyeh K Chu, and Jerome Mertz, “Wide-field fluorescence sectioning with hybrid speckle and uniform-illumination microscopy,” Opt. letters 33, 1819–1821 (2008).

157. Daryl Lim, Timothy N Ford, Kengyeh K Chu, and Jerome Metz, “Optically sectioned in vivo imaging with speckle illumination HiLo microscopy,” J. biomedical optics 16, 016014 (2011).

158. Tyler D Ross, Heun Jin Lee, Zĳie Qu, Rachel A Banks, Rob Phillips, and Matt Thomson, “Controlling organization and forces in active matter through optically defined boundaries,” Nature 572, 224–229 (2019).

159. Pavel Křížek, Ivan Raška, and Guy M Hagen, “Flexible structured illumination microscope with a programmable illumination array,” Opt. express 20, 24585–24599 (2012).

160. Ziwei Li, Qinrong Zhang, Shih-Wei Chou, Zachary Newman, Raphaël Turcotte, Ryan Natan, Qionghai Dai, Ehud Y Isacoff, and Na Ji, “Fast widefield imaging of neuronal structure and function with optical sectioning in vivo,” Sci. Adv. 6, eaaz3870 (2020).

161. Hui-Wen Lu-Walther, Martin Kielhorn, Ronny Förster, Aurélie Jost, Kai Wicker, and Rainer Heintzmann, “fastSIM: a practical implementation of fast structured illumination microscopy,” Methods Appl. Fluoresc. 3, 014001 (2015).

162. Alice Sandmeyer, Mario Lachetta, Hauke Sandmeyer, Wolfgang Hübner, Thomas Huser, and Marcel Müller, “DMD-based super-resolution structured illumination microscopy visualizes live cell dynamics at high speed and low cost,” bioRxiv p. 797670 (2019).

163. Meiqi Li, Yaning Li, Wenhui Liu, Amit Lal, Shan Jiang, Dayong Jin, Houpu Yang, Shu Wang, Karl Zhanghao, and Peng Xi, “Structured illumination microscopy using digital micro-mirror device and coherent light source,” Appl. Phys. Lett. 116, 233702 (2020).

164. Peter T Brown, Rory Kruithoff, Gregory J Seedorf, and Douglas P Shepherd, “Multicolor structured illumination microscopy and quantitative control of coherent light with a digital mirror device,” bioRxiv (2020).

165. Howard C. Berg, “How to track bacteria,” Rev. Sci. Instruments 42, 868–871 (1971).

166. Alan Lukežic, Tomáš Vojíř, Luka Čehovin Zajc, Ji vr’i Jiří Matas, Matej Kristan, Tom’aš Voj’i vr, Luka Čehovin Zajc Ji vr’i, Jiří Matas, and Matej Kristan, “Discriminative Correlation Filter Tracker with Channel and Spatial Reliability,” Int. J. Comput. Vis. 126, 671–688 (2018).

167. Eugen Baumgart and Ulrich Kubitscheck, “Scanned light sheet microscopy with confocal slit detection,” Opt. express 20, 21805–21814 (2012).

168. Raju Tomer, Li Ye, Brian Hsueh, and Karl Deisseroth, “Advanced CLARITY for rapid and high-resolution imaging of intact tissues,” Nat. protocols 9, 1682 (2014).

169. MA Model and JL Blank, “Concentrated dyes as a source of two-dimensional fluorescent field for characterization of a confocal microscope,” J. microscopy 229, 12–16 (2008).

170. Kyle M Douglass, Christian Sieben, Anna Archetti, Ambroise Lambert, and Suliana Manley, “Super-resolution imaging of multiple cells by optimized flat-field epi-illumination,” Nat. photonics 10, 705–708 (2016).

171. Christian M Schürch, Salil S Bhate, Graham L Barlow, Darci J Phillips, Luca Noti, Inti Zlobec, Pauline Chu, Sarah Black, Janos Demeter, David R McIlwain et al., “Coordinated cellular neighborhoods orchestrate antitumoral immunity at the colorectal cancer invasive front,” Cell 182, 1341–1359 (2020).

172. Hagai Kirshner, Franois Aguet, Daniel Sage, and Michael Unser, “3-D PSF fitting for fluorescence microscopy: implementation and localization application,” J. microscopy 249, 13–25 (2013).

173. Zheng Zhu, Qiang Wang, Li Bo, Wei Wu, Junjie Yan, and Weiming Hu, “Distractor-aware Siamese Networks for Visual Object Tracking,” in European Conference on Computer Vision, (2018).

